# Regulation by partition predicts enhancer target selection in mammalian genomes

**DOI:** 10.64898/2026.10.23.761042

**Authors:** Jordan Valgardson, Tomek Swigut, Joanna Wysocka

## Abstract

Enhancers regulate specific promoters across long genomic ranges, yet how they select among the many promoters within reach remains unresolved. Here we introduce SPIE-seq (Synthetic Perturbation by Inserted Enhancer), which pairs piggyBac-mediated enhancer insertion with an allele-specific readout of cis-regulatory effects on expression. Across 724 clonal lines carrying 29,954 mapped insertions, we detected enhancer-dependent expression changes at 18,440 enhancer-gene pairs. We found that enhancer influence declines with genomic distance, following similar average distance-decay scaling across all tested enhancers, though the shape of response curves differs between loci. Genomic context, not enhancer identity, dominates target choice: distinct enhancers inserted at the equivalent positions activate nearly the same targets, while the same enhancer can activate discordant promoters when inserted at different positions within the same locus. Responsive promoters are enriched for developmental features, but target choice is nonetheless competitive: even at loci with high promoter density, an enhancer primarily activates only one or two promoters. Promoter skipping is common and increases as competitors accumulate — a pattern incompatible with promoters responding independently based solely on their intrinsic features. We formalize this in the Regulation-by-Partition (RbP) model, in which an enhancer allocates its output among the promoters according to their genomic distance, contact with the enhancer, and intrinsic features. RbP predicts target choice accurately and transfers without retraining across species, cell types, and perturbation direction, outperforming existing enhancer–gene models, with the largest gains at long genomic ranges. Enhancer target choice is thus not dictated by enhancer or promoter attributes alone but emerges from the distance-weighted competition among promoters within the regulatory landscape.

## Introduction

Enhancers are the principal long-range cis-regulatory elements of metazoan genomes that act by increasing transcription of a target promoter, often from distances of tens of kilobases of intervening DNA and, in extreme cases, more than a megabase (*1–4*). This reach creates a fundamental puzzle: any megabase-scale window in the genome typically contains multiple candidate promoters, yet enhancers reliably regulate specific targets while ignoring others. How enhancers achieve this specificity across such large genomic distances remains a central open question in gene regulation.

Strong evidence for enhancer-promoter specificity comes from endogenous loci where enhancers demonstrably bypass intervening promoters — a phenomenon known as promoter skipping. For example, at the β-globin locus, the locus control region (LCR) bypasses the more proximal embryonic and fetal globin promoters to selectively activate the adult β-globin gene (HBB) in adult erythroid cells (*5–7*). Skipping of this kind is difficult to reconcile with a null model in which enhancers indiscriminately activate any promoter within contact range. Whether skipping reflects sequence-encoded enhancer–promoter compatibility, intrinsic features of individual promoters, or emergent properties of the local genomic landscape remains unclear.

One proposed determinant of enhancer target selection is the intrinsic regulatory character of the promoter itself. Metazoan promoters have long been divided into two broad categories: housekeeping promoters, which drive genes expressed broadly across cell types with low variability, and developmental promoters, which drive cell type-restricted genes whose expression changes markedly during differentiation. Recent work has proposed that housekeeping promoters operate near the saturation of their transcriptional output and are only weakly responsive to boosting by enhancers, whereas developmental promoters retain the dynamic range required to respond (*8–10*). The properties underlying this distinction, however, are continuous: various measures of expression breadth and variability across cell types, as well as promoter architecture, place promoters along a gradient rather than partitioning them cleanly (*11–15*). How such intrinsic properties combine with the arrangement of regulatory elements in the local landscape to determine enhancer targets remains untested.

Progress in identifying principles that govern enhancer target choice has been constrained by the available large-scale methods for probing how (and which) promoters respond to enhancers in the genomic environment. Massively parallel reporter assays (MPRAs) scale well but strip enhancers from their genomic context. Consequently, these assays – although extremely useful for inferring autonomous regulatory potential of enhancers and promoter boostability – have largely failed to recapitulate the enhancer–promoter specificity observed at endogenous loci (*16*, *17*). Loss-of-function perturbations — enhancer deletions and CRISPRi-mediated silencing — provide direct evidence of regulatory dependencies but remain limited in two important respects. First, scaling these approaches across tens of thousands of candidate enhancers active in a given cell type remains challenging (*16*, *18*, *19*), and second, since removal-based strategies interrogate enhancers one at a time within complex regulatory landscapes, redundant enhancers can mask the contribution of any single element, making absence of a detectable effect difficult to interpret (*20*).

A complementary strategy is to ask not what happens when an enhancer is removed from its native locus, but what happens when a well-characterized enhancer is introduced into diverse contexts across the genome. This bottom-up approach — adding a single, defined regulatory element and measuring its effects on all surrounding promoters — offers several advantages. Since the inserted enhancer is a gain-of-function perturbation, its effects are not subject to masking by redundant elements. Because the same enhancer lands in thousands of unrelated genomic neighborhoods through random integration, and because different enhancers can land in the same neighborhood, the approach naturally separates enhancer identity, genomic distance, local promoter composition, and chromatin environment — variables that are intertwined at any single endogenous locus and difficult to disentangle even across large perturbational datasets. Random integration also makes this tractable at scale, providing the statistical power that quantitative modeling of enhancer target choice requires.

Here, we describe SPIE-seq (Synthetic Perturbation by Inserted Enhancer), an assay that combines piggyBac-mediated insertion of well-characterized enhancers in F1 hybrid mouse embryonic stem cells (mESC) with allele-specific readout of cis-regulatory expression effects. Across 724 clonal mESC lines carrying 29,954 mapped insertions, we detected enhancer-dependent expression changes at 18,440 enhancer–gene pairs spanning 9,143 distinct promoters, thus representing the most comprehensive enhancer perturbation dataset generated to date. We found that enhancer influence on gene expression is approximately constant below ∼10 kb and then decays steeply with genomic distance, with the short-range plateau consistent with the saturation of the promoter’s output. Enhancers of different identities followed similar distance-dependent scaling on average, but the shape of this relationship varied between loci depending on the target promoter’s dynamic range and the promoter density within the locus. Moreover, when different enhancers are inserted at the same position, they activate nearly the same promoter targets, indicating that the local genomic landscape, not the enhancer identity, is the dominant determinant of target choice.

Enhancers frequently act beyond the nearest promoter: in ∼28% of cases activation extends to a promoter other than the closest one, and in ∼18% of cases — a subset of these — the enhancer skips over at least one intervening promoter without activating it. Responsive promoters, especially those activated at long-range, are enriched for features associated with developmental promoters, including higher expression dynamic range and expression inequality across tissues, as well as enrichment of H3K27me3 and lower level of H3K27ac; these features are continuous rather than binary, but place responsive promoters toward the developmental end of the spectrum. However, intrinsic promoter attributes and distance are insufficient to explain target choice. Comparing insertions at independent positions near the same promoter, we find that a promoter can respond to the enhancer placed further away and fail to respond when the same enhancer is placed closer but other promoters lie nearer still. Overall, an enhancer activates 1.35 promoters on average, and this number rises only marginally with the number of promoters within reach, even at loci containing tens of them. Promoters within a regulatory landscape thus compete for a limited enhancer output, behavior incompatible with promoters responding independently according to their own features. We formalize these observations in the Regulation-by-Partition (RbP) model, in which each promoter receives a score from its genomic distance, predicted contact with the enhancer and intrinsic features, while competition is encoded by partitioning the finite enhancer output among promoters, in proportion to their scores. The RbP outperforms an unconstrained gradient-boosted model trained on a large set of features despite using far fewer parameters and predicts enhancer target choice across our dataset with high accuracy. We further show that the RbP model outperforms existing state of the art enhancer–gene models, such as rE2G and AlphaGenome, in predicting, without retraining, the outcomes of CRISPRi enhancer silencing on expression in human K562 cells (*18*, *21–23*). Superior performance of RbP over other existing models is especially evident for regulatory interactions that occur at long genomic ranges of over 100 kb. Finally, the RbP model accurately predicts regulatory effects of enhancer deletions in mESC, of promoter deletions at the β-globin locus in erythroid cells, and of eQTL SNPs in human fibroblasts. Thus, the RbP generalizes across cell types, species, and direction of enhancer perturbation, suggesting that the quantitative logic for enhancer target selection described here captures key features of mammalian long-range gene regulation.

## Results

### SPIE-seq: Synthetic Perturbation by Inserted Enhancer

To systematically interrogate promoter responsiveness to enhancers across diverse genomic contexts, we developed Synthetic Perturbation by Inserted Enhancer (SPIE) seq, which measures the transcriptional consequences of enhancer insertions at scale. In SPIE-seq, enhancer sequences are inserted at random genomic positions via piggyBac transposase and clonal mESC lines are isolated (Fig. 1A). We optimized transposition conditions such that each cell line contains 10-100 insertions – this enables scoring multiple genomic contexts per cell line, while keeping the overall insert density in the genome low (estimated 1 per 27 Mb at 100 insertions). Insert locations are mapped by splinkerette sequencing, and the transcriptional effects of each insertion are measured by RNA-seq (*24*). To enable allele-specific quantification of enhancer effects in cis, while controlling for potential trans effects, we performed SPIE-seq in the 129×CAST F1 hybrid mouse embryonic stem cell (mESC) line F121. Insertions occur on one allele, while the other serves as an internal control (Fig. 1A).

**Figure 1.**
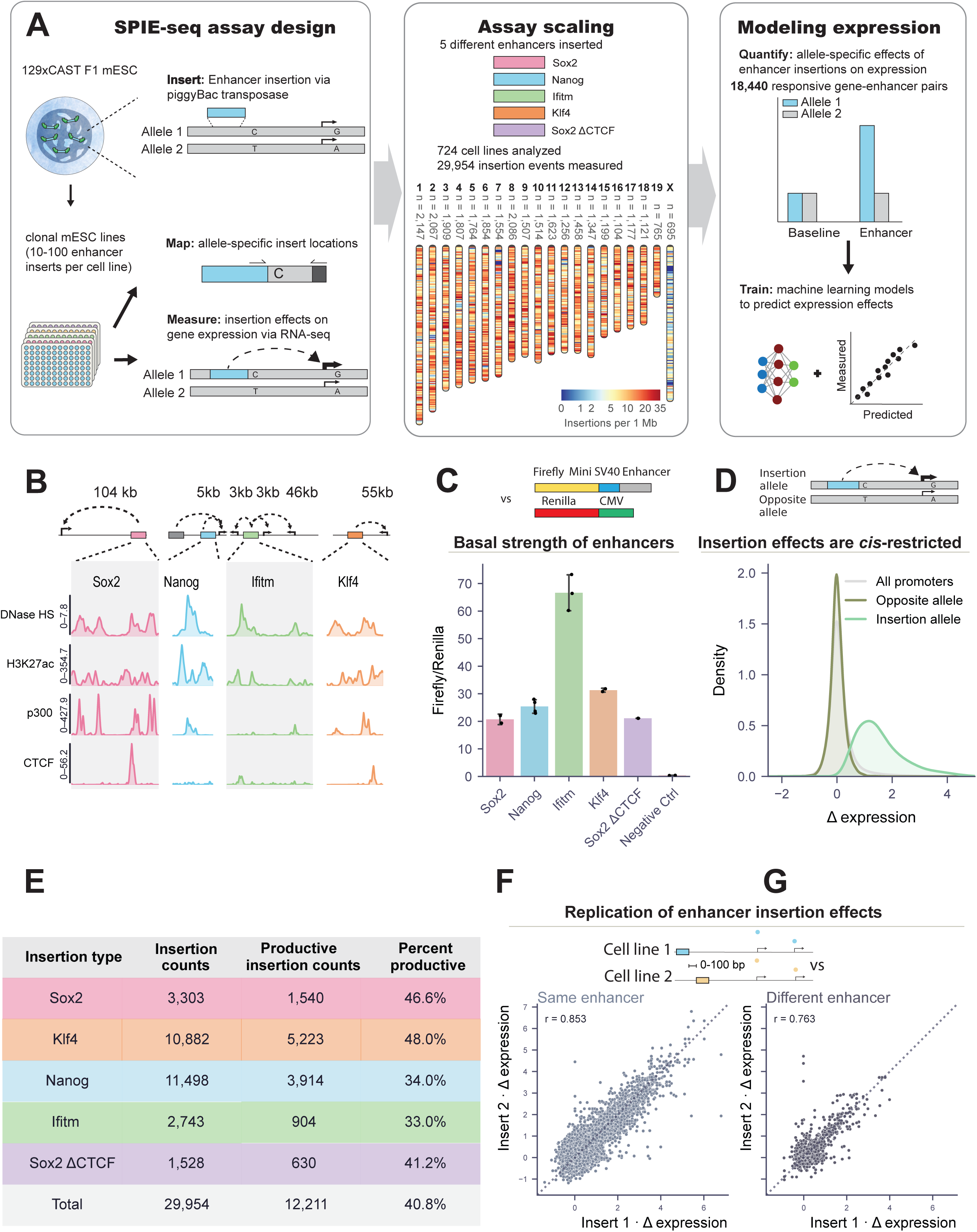
SPIE-seq maps allele-resolved cis-regulatory effects of enhancer insertions across thousands of genomic contexts and reveals that genomic position, not enhancer identity, determines target promoter choice. **(A)** SPIE-seq assay design. Enhancers were inserted at random genomic positions by piggyBac transposase in 129 × CAST F1 hybrid mESC and clonal lines with 10-100 inserts per cell were isolated. Insert positions were mapped by splinkerette sequencing and allele-assigned using strain-specific SNPs; expression was measured by allele-resolved RNA-seq. Ideogram: insertion density in 1-Mb bins; n, total insertions per chromosome. **(B)**Native chromatin environment of each inserted enhancer. Top: locus schematics showing the inserted fragment (colored), other enhancers at the locus (grey), promoters (arrows) and native enhancer–promoter distances (dashed arrows). Bottom: wild-type mESC DNase hypersensitivity (read-normalized signal), H3K27ac and p300 ChIP-seq (signal p-value), and CTCF ChIP-seq (spike-in-normalized coverage) over each fragment. **(C)** Luciferase reporter activity. Each indicated enhancer was cloned upstream of a minimal SV40 promoter driving firefly luciferase, and co-transfected with CMV-Renilla into mESC. Negative control: SV40 promoter alone. Bars, firefly/Renilla ratio; points, three replicates; error bars, s.d. **(D)** Cis-specificity. Δ expression (shrunk log fold change) at allele-resolved insertion sites. Promoters within 200 kb responding on the insertion-bearing allele (green), the same promoters on the homologous allele (dark green), and all promoters genome-wide (grey). **(E)** Number of insertions and productive insertions per enhancer construct. An insertion is productive if it upregulates at least one promoter within 500 kb (FDR < 5%; see Methods). **(F)** Reproducibility. Δ expression for independently derived lines carrying the same enhancer within 100 bp of each other (28,844 comparisons, 1,966 site pairs; r = 0.853). Dashed line, y=x. (G) As in (F), for different enhancers inserted within 100 bp of each other (2,002 comparisons, 142 site pairs; r = 0.763).

We selected four distinct enhancers for insertion based on well-established genetic evidence of their major contribution to gene expression in mESC (Fig. 1B). The *Sox2* control region (SCR) is a well-characterized enhancer cluster located ∼100 kb from the *Sox2* promoter; deletion of the SCR reduces *Sox2* expression to approximately 10% of wild-type levels (*25*, *26*). The *Klf4* enhancer, located ∼55-68 kb from *Klf4*, drives the majority of *Klf4* expression (*27*). The *Ifitm* enhancer resides in the *Ifitm* locus and drives expression of *Ifitm1*, *Ifitm2* and *Ifitm3* (*28*). The *Nanog* enhancer, located 5 kb upstream of the *Nanog* promoter, is responsible for nearly all *Nanog* expression (*29*). Finally, we generated a variant of the *Sox2* enhancer in which all CTCF binding sites—both occupied and unoccupied—were removed (*Sox2* ΔCTCF). We included this construct to test whether CTCF sites within an enhancer influence its long-range promoter selection at ectopic loci. Deleting these sites at the endogenous locus has minimal effect on *Sox2* expression (*25*), and we sought to determine whether this held true across diverse genomic contexts.

To compare intrinsic transcriptional activating capacity of these five enhancer constructs, we tested each in a luciferase reporter assay with an SV40 minimal promoter. All five constructs boosted luciferase expression above the negative control, with the *Ifitm* enhancer showing the strongest activity (Fig. 1C). Importantly, the wild-type and *Sox2* ΔCTCF enhancers produced equivalent luciferase output, confirming that removing CTCF sites does not alter the enhancer’s intrinsic activation strength (Fig. 1C).

In total, we generated 724 clonal mESC lines harboring enhancer insertions, yielding 29,954 mapped insertion events across the genome, with robust representation of each of the five enhancers (Fig. 1A, E). To quantify transcriptional effects measured by RNA-seq across this cell line panel, we developed a custom differential expression framework adapted to the single-replicate structure of the SPIE-seq data. Briefly, for each insert we consider all promoters within 2 Mb of the insertion. For every promoter, we estimate a background distribution from all cell lines that did not receive an insert within 2 Mb, and use it to compute an empirical p-value for the expression measured in lines carrying a proximal insert. We apply FDR correction by estimating the false-positive rate in regions far from any insert (>2 Mb), and we call a promoter regulated when its false discovery rate for upregulation falls below 5 percent (see Methods for details).

Of the 29,954 insertion events, 40.8% upregulated at least one promoter within 500 kb (Fig. 1E). This productive fraction may appear modest, but many insertions land in large gene deserts with no nearby promoters. In addition, we cannot rule out the possibility of silencing of the inserts in some genomic contexts. Nonetheless, we scored over twelve thousand ‘productive’ insertion events, where at least one promoter changed expression in cis upon enhancer insertion (Fig. 1E). Across the dataset, we scored 18,440 significant enhancer– promoter pairs, involving 9,143 distinct promoters, with many promoters responding to insertions in more than one cell line. Thus, our SPIE-seq screen captured enhancer responses across a large number of regulatory environments in the mouse genome.

To test whether inserted enhancers could act in trans, we leveraged the hybrid nature of the F121 line. For each insertion we identified promoters within 200 kb that responded significantly on the insert-bearing allele, then examined the same promoters on the homologous allele. Effects were restricted to the allele bearing the insertion (Fig. 1D): the median effect on the homologous allele was indistinguishable from zero, and the mean residual was under 5% of the cis effect — within the range expected from imperfect allelic read assignment. We therefore do not interpret this residual as evidence of trans activity, consistent with the expectation that transvection of the kind described in *Drosophila* (*30–32*) is rare in mammalian cells, and supporting the use of the non-insert allele as an internal control.

### Genomic context, rather than enhancer identity, shapes promoter responses

While resequencing all 724 cell lines was prohibitively expensive, the high density of insertions across our panel afforded a more stringent measure of assay reproducibility: we identified cases in which independent clonal lines received insertions at the equivalent genomic location. Because these lines are independent clones, this comparison decouples clone-specific trans effects from the local activation being measured. When the same enhancer was inserted within 100 bp in two independent lines, the transcriptional effects on nearby promoters were highly correlated (Pearson r = 0.853, n = 1,966 regions with two insertions; Fig. 1F), attesting to high reproducibility of our results across different cell lines and independent insertion events. Strikingly, when different enhancers were inserted at the same location, the correlation remained strong (r = 0.763, n = 142 regions with two insertions; Fig. 1G), suggesting that genomic context—rather than enhancer identity—is the dominant determinant of which promoters respond, and how much.

### Expression responses decay with E-P distance similarly across enhancers and are restricted by TAD boundaries

Across all insertion events, both the magnitude and the probability of activation fell with the distance between the insertion and the promoter, and insertions rarely repressed nearby promoters (Fig. 2A). This is consistent with previous reports of enhancer activity decay with distance from the promoter (*33–38*). To characterize average distance-decay profiles and compare them between different enhancers, we calculated aggregate decay fits across insertion–promoter pairs within 1 Mb of an isolated insertion, in regions in which at least one promoter responds within 500 kb. For all enhancers, the relationship is not a simple power law: response is approximately constant below ∼10 kb, decays over the following decade of distance, and approaches zero at 1 Mb (R² = 0.92–0.97 per enhancer, 0.99 pooled, on binned means; Fig. 2B). Curves for the Sox2 enhancer with and without CTCF sites were similar, indicating that enhancer-internal CTCF sites affect neither the range nor the shape of distance decay.

**Figure 2.**
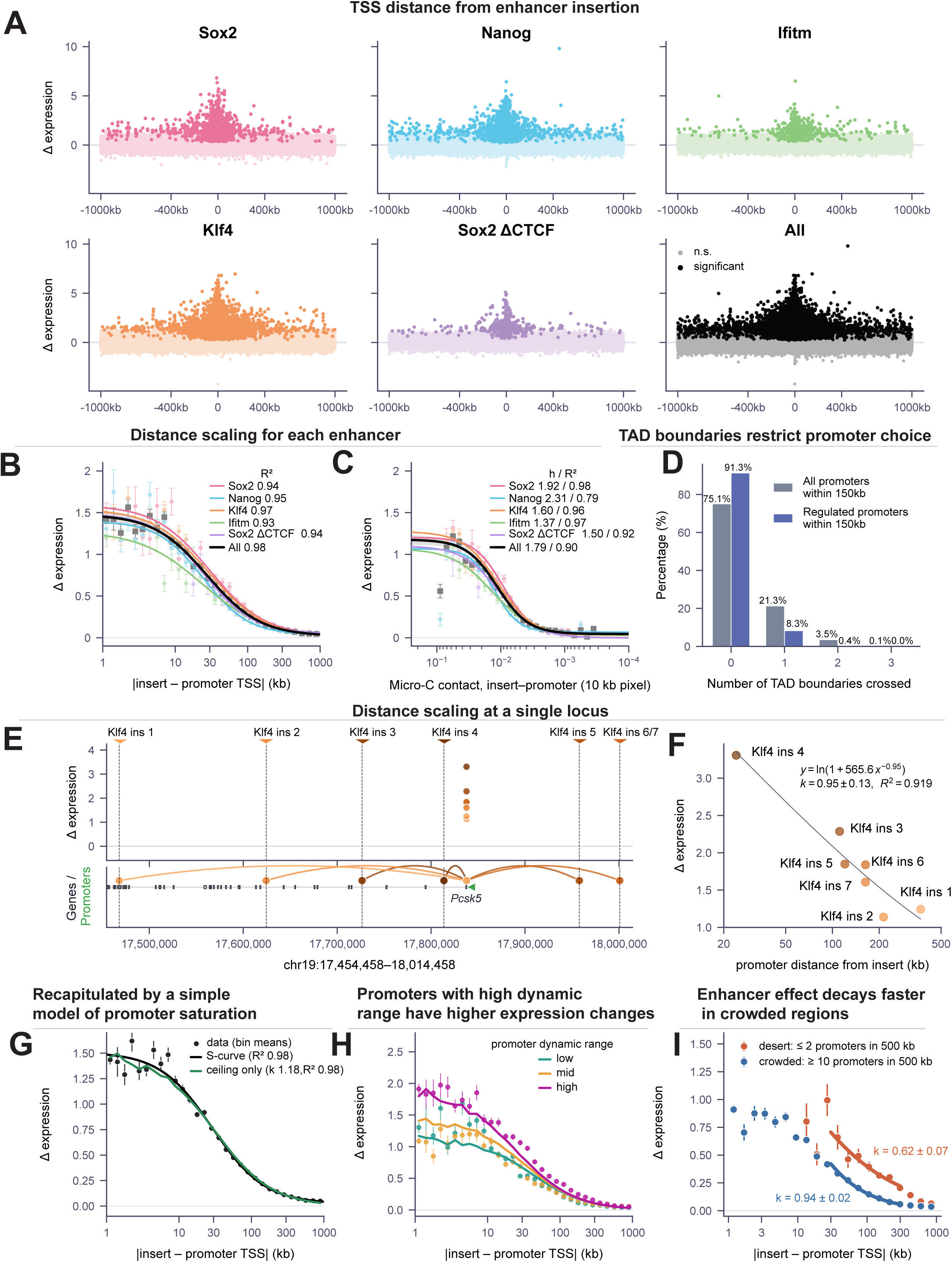
Enhancer activity follows an S-shaped decay with genomic distance consistent with promoter saturation, and is modulated by 3D contact, TAD boundaries, and local gene density. **(A)** Effect of each insertion on all promoters within 1 Mb, by construct. Δ expression (shrunk log fold change) versus signed insert–TSS distance. Dark points, significantly upregulated (ln p-value < −10); pale, not significant. **(B)** Average enhancer effect as a function of enhancer–promoter distance. Mean Δ expression (shrunk log fold change) of enhancer–promoter pairs versus absolute insert–TSS distance for pairs in (A). Points, means of logarithmic bins (30 logarithmically spaced bins; bins with <20 promoters not drawn); error bars, ±s.e.m. Lines, four-parameter logistic fits; R² per construct in key. Black band (pooled curve), 95% percentile interval from 200 bootstrap refits resampling insertions. **(C)** Average enhancer effect as a function of 3D contact. Mean Δ expression (shrunk log fold change) versus Micro-C contact frequency (10 kb pixel; mESC data from ref. *39*). Pairs with zero contact excluded. Lines, Hill fits; key gives Hill coefficient (h) and R². Black band, 95% bootstrap interval. **(D)** Depletion of enhancer–promoter communication across TAD boundaries. Percentage of all promoters (grey) and significantly regulated promoters (blue) at each boundary count, for promoters within 150 kb of an insertion (no other insertion within 1 Mb; ref. *44*). n = 100,581 total pairs; 4,631 significant pairs. **(E)** Distance-dependent activation at a single locus. Seven insertions of the klf4 enhancer across six positions across the *Pcsk5* locus in independent clonal lines. Top, Δ expression (shrunk log fold change) per promoter; point color, insertion distance from the *Pcsk5* promoter; dashed lines and triangles, insertion sites. Bottom, gene models with arcs connecting significantly upregulated enhancer–promoter pairs (ln p-value < −10). **(F)** Power-law decay at the *Pcsk5* locus. Δ expression at *Pcsk5* versus distance from the inserted Klf4 enhancer (n = 7 independent lines). Grey line, fit of a saturating power law, Δ = ln(1 + b·d^-k), in which the effect decays as an inverse power of distance and saturates at short range; k = 0.95 ± 0.13, R² = 0.919. **(G)** A promoter-capacity model matches the aggregate distance–response curve. Points, bin means ± s.e.m. over all enhancer-promoter pairs (n = 141,850; 30 logarithmically spaced bins). Black, four-parameter S-curve (R² = 0.98). Green, ceiling-only model: power-law decay (exponent k = 1.18) capped by each promoter’s dynamic range (5th–95th percentile of FANTOM5 CAGE expression), R² = 0.98. **(H)** Promoters with more dynamic range respond more strongly at short distances. Bin means ± s.e.m. of Δ expression (shrunk log fold change) versus distance, with promoters split into terciles of dynamic range ranked within each distance bin (∼47,280 promoters per tercile). Lines, ceiling-model fits. **(I)** Enhancer effects reach farther in gene-sparse regions. Bin means ± s.e.m. of Δ expression (shrunk log fold change) versus distance for insertions in gene deserts (≤2 promoters within 500 kb; orange) versus crowded regions (≥10 promoters within 500 kb; blue). Solid lines, fits of a saturating power law beyond 20kb; k ± s.e. as shown.

Interestingly, the same S-shape response holds when expression changes are compared against chromatin contact. Using published mESC Micro-C (*39*), we estimated contact between each insertion’s flanking region and each promoter and fit a Hill curve of response against contact: response rises from near zero at low contact to a plateau at high contact, with Hill coefficients of 1.37–2.31 across the five enhancers (Fig. 2C).

### TAD boundaries restrict promoter choice beyond what distance and contact predict

While genomic distance is the dominant predictor, TAD boundaries impose an additional constraint (*40–43*). Using TAD definitions from Bonev et al. 2017 (ref. *44*), and restricting to promoters within 150 kb of an insertion to control for the confounding of distance with TAD size, 21.3% of promoters lie across at least one boundary, yet only 8.3% of the promoters actually activated by the enhancer do so — an approximately 3-fold reduction in activation probability (Fig. 2D;*44*). The depletion holds at every distance cutoff from 50 kb to 500 kb (Sup. Fig. 1A). TAD boundaries therefore restrict, but do not abolish, enhancer–promoter communication, consistent with other recent studies (*45–49*).

### Promoter response ceiling explains locus-specific distance scaling relationships

Previous studies at individual loci reported inconsistent distance scaling relationships, with some studies describing a scale-free power law, others a curve that is flat at short range and falls steeply thereafter (*33–38*). Indeed, in our dataset we observed that despite the global S-shape distance response across all enhancers, loci differed in shape of their distance responses. For example, the *Pcsk5* locus received seven independent Klf4 enhancer insertions within 500 kb across different clonal lines, none of which carried additional nearby insertions that could confound the analysis. *Pcsk5* expression decays with enhancer distance according to a power-law with an exponent of 0.95 ± 0.13 (R² = 0.92; Fig. 2E, F). The nearest Klf4 insertion is located >20 kb from the *Pcsk5* promoter, which may be beyond the plateau observed in Fig. 2B. Nonetheless, our observations point to a fundamental tension: enhancer activity scaling behavior appears to differ between promoters.

We hypothesized that the plateau at short range may be associated with saturation: once an enhancer is close enough, bringing it closer does not increase output, because the promoter reaches the ceiling of its response. Thus, which loci show the S-shape vs power-law scaling may depend on how much headroom for activation the promoter has before it saturates. To explore this idea further, we tested whether a minimal model — a power-law decay in distance combined with a promoter saturation ceiling set by the promoter’s dynamic range between the 5th and 95th percentiles of that promoter’s expression across the FANTOM5 CAGE-seq dataset — could reproduce the observed curve. It does (power exponent 1.18, R² = 0.98 on binned means; Fig. 2G). A four-parameter empirical S-curve fits just as well (R² = 0.98).

The ceiling also predicts which promoters respond most. Stratifying by dynamic range separates the response curves vertically; high dynamic range promoters show the largest responses and low dynamic range promoters the smallest (Fig. 2H). Headroom then determines how power-law-like the decay appears. The decay follows a power law wherever the response sits below the ceiling and is flat wherever the response saturates the promoter, so the apparent shape over any measured window depends on how much headroom the promoter has. Promoters with ample headroom are rarely saturated and scale as a power law across most of the accessible distance range; promoters near their ceiling are saturated out to longer distances and appear S-shaped. This may reconcile the locus-dependent differences in distance scaling of enhancer activity reported in the literature (*33*, *34*, *36–38*, *42*).

Moreover, we observed that the scaling behavior of enhancers depends on gene density. We aggregated, without further conditioning, regions with two or fewer promoters within 500 kb of the enhancer (gene deserts) and regions with ten or more (crowded loci). In gene deserts, responses are larger and decay more shallowly beyond ∼20 kb than in crowded loci (fitted exponents k = −0.62 ± 0.07 versus −0.94 ± 0.02;Fig. 2I). Taken together, these results imply that enhancer scaling laws are locus-specific, set by promoter saturation and gene density.

### Enhancers frequently skip the nearest promoter

Having established that genomic distance is a key determinant of enhancer regulatory reach, we turned to promoter skipping—instances in which an enhancer activates a more distal promoter while leaving a closer one unaffected. This phenomenon is well-documented in the literature (*1*, *5*, *6*, *35*, *50*, *51*), but our unique dataset allows a genome-wide estimate of its frequency. Across the dataset, considering only insertions that activated at least one promoter within 500 kb, the enhancer activated the nearest promoter (rank distance = 1, counted from the side of insertion) approximately 72% of the time (Fig. 3A). In the remaining ∼28% of cases the activated target was the second-nearest promoter or further. Not all of these represent true skipping, since an enhancer may activate a distal promoter while also activating the nearer one; restricting to cases in which the intervening promoter was left unaffected, enhancers bypassed at least one promoter entirely in ∼18% of cases (Sup. Fig. 2A). Skipping was observed for all enhancers tested, though the fraction of insertions targeting the nearest promoter varied from ∼68% for the Ifitm enhancer to ∼84% for the Nanog enhancer (Fig. 3A). An illustrative example occurs at the *Mycl* locus, where a Nanog enhancer inserted in an intron of Trit1 bypasses Trit1, an expressed gene, to upregulate *Mycl* 100 kb away (Fig. 3B). Altogether, promoter skipping by enhancers is relatively frequent across the genome.

**Figure 3.**
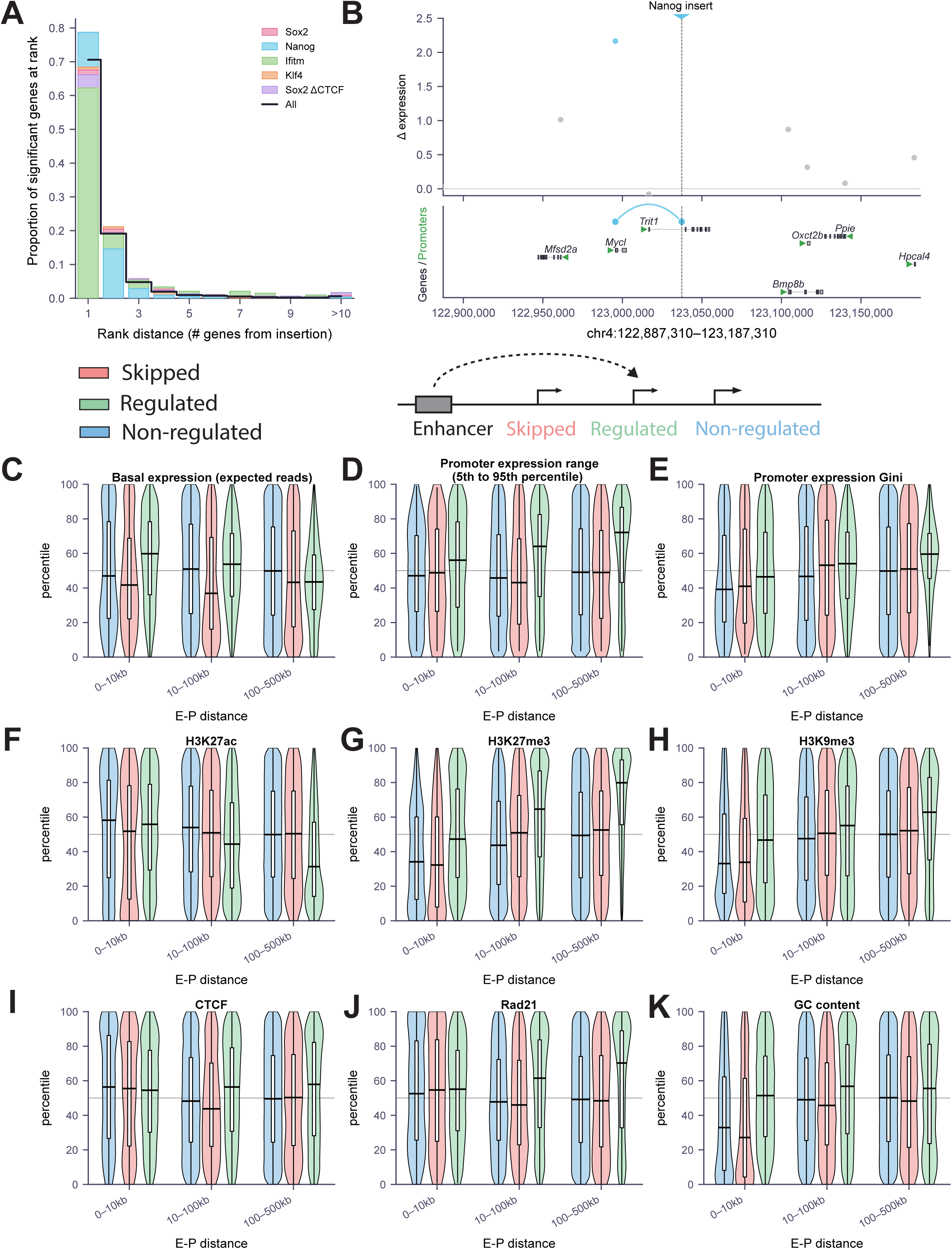
Enhancers frequently skip proximal promoters, and long-range responsive promoters are enriched for developmental features. **(A)** Distribution of rank distance between an insertion and the promoters it significantly upregulates. Rank distance is the position of the activated promoter among all promoters on that side of the insertion (rank 1, nearest). **(B)** Example: Nanog enhancer inserted into an intron of *Trit1* (chr4:122,887,310– 123,187,310). Top, Δ expression (shrunk log fold change) per promoter; blue, significantly upregulated; grey, not significant; dashed line and triangle, insertion site. Bottom, gene models with arcs connecting significant enhancer–promoter pairs. **Schematic (above C).** Promoter classes: regulated (green), significantly upregulated; skipped (pink), not upregulated but lying between the insertion and a regulated promoter; non-regulated (blue), neither but within 500 kb of the enhancer. **(C–K)** Genomic and epigenomic features of non-regulated, skipped and regulated promoters, stratified by enhancer–promoter distance (0–10 kb, 10–100 kb, 100–500 kb). Each feature expressed as a percentile rank across all promoters; grey line, 50th percentile. **(C)** Basal expression; **(D)** expression range (FANTOM5, p95 − p5); **(E)** expression Gini coefficient; **(F)** H3K27ac; **(G)** H3K27me3; **(H)** H3K9me3; **(I)** CTCF; **(J)** Rad21; **(K)** GC content. Violins with overlaid box-and-whisker plots (median, IQR, 1.5× IQR whiskers).

### Promoter features associated with enhancer responsiveness

What determines whether an enhancer skips or regulates a proximal promoter? To address this, we compared 1,151 genomic and epigenomic features across three classes of promoters: those activated by a nearby insertion (“regulated”), those closer to the enhancer but not activated (“skipped”), and those outside the enhancer’s apparent regulatory range (“non-regulated”) (see schematics in Fig. 3; all analyzed promoter features are listed in Supplementary Table 1). Of 1,151 promoter features, we retained 173 after removing uninformative features, and prioritized those with the highest importance in a gradient-boosted classifier, trained to distinguish regulated from skipped and non-regulated promoters. The most predictive features fall into four groups: measures of nascent transcription and transcript metabolism in mESC (GRO-seq signal, RNA synthesis rate from 5-EU pulse-chase, and transcript length); H3K27me3 enrichment at the promoter; the promoter’s dynamic range of expression across tissues (5^th^-95^th^ percentile range and interquartile range); and binding of cohesin (Rad21, Smc1A) and other architectural/coactivator proteins (Ldb1, Yy1) at the promoter (Sup. Fig. 2B). We have focused our subsequent analyses on these features and a few others that have been previously associated with promoter activity/silencing and long-range contacts (e.g. H3K27ac, H3K9me3, GC content and CTCF binding). Surprisingly, even for the most discerning features, their marginal distributions overlapped substantially across all three categories (Sup. Fig. 3).

We explored expression features of the three promoter classes in their unperturbed state (that is, without the ectopic enhancer insertion), analyzing each feature as a function of distance from the enhancer insertion by separating each promoter class into three bins based on the E-P distance. All three classes showed broad and overlapping basal expression range, indicating that simple ‘active/inactive’ classification cannot explain enhancer responsiveness (Fig. 3C). Interestingly, however, the association between basal expression level and promoter responsiveness becomes more evident at greater distances, where genes driven by regulated promoters show lower median basal expression than those driven by either skipped or non-regulated promoters (Fig. 3C).

We considered whether the association between low basal expression and responsiveness at long ranges might reflect detection sensitivity: a fixed absolute increase in transcription produces a larger fold change — and thus a more readily detected signal — at a lowly expressed promoter than at a highly expressed one. Our data argue against this. Because expression is quantified from read counts, statistical power is governed by counting noise, which is proportionally largest at low expression. The minimum fold change required to call a significant change is therefore steepest for weakly expressed promoters, requiring 5- to 17-fold induction in the lowest expression bins, compared with under 2-fold for the most highly expressed promoters (Sup. Fig. 4A). Consistent with this, significantly regulated promoters are enriched, not depleted, among promoters with higher expected read counts relative to all promoters within 100 kb of an insertion (Sup. Fig. 4B). The enrichment of lowly expressed promoters among long-range targets therefore occurs despite, rather than because of, the detection properties of the assay.

**Figure 4.**
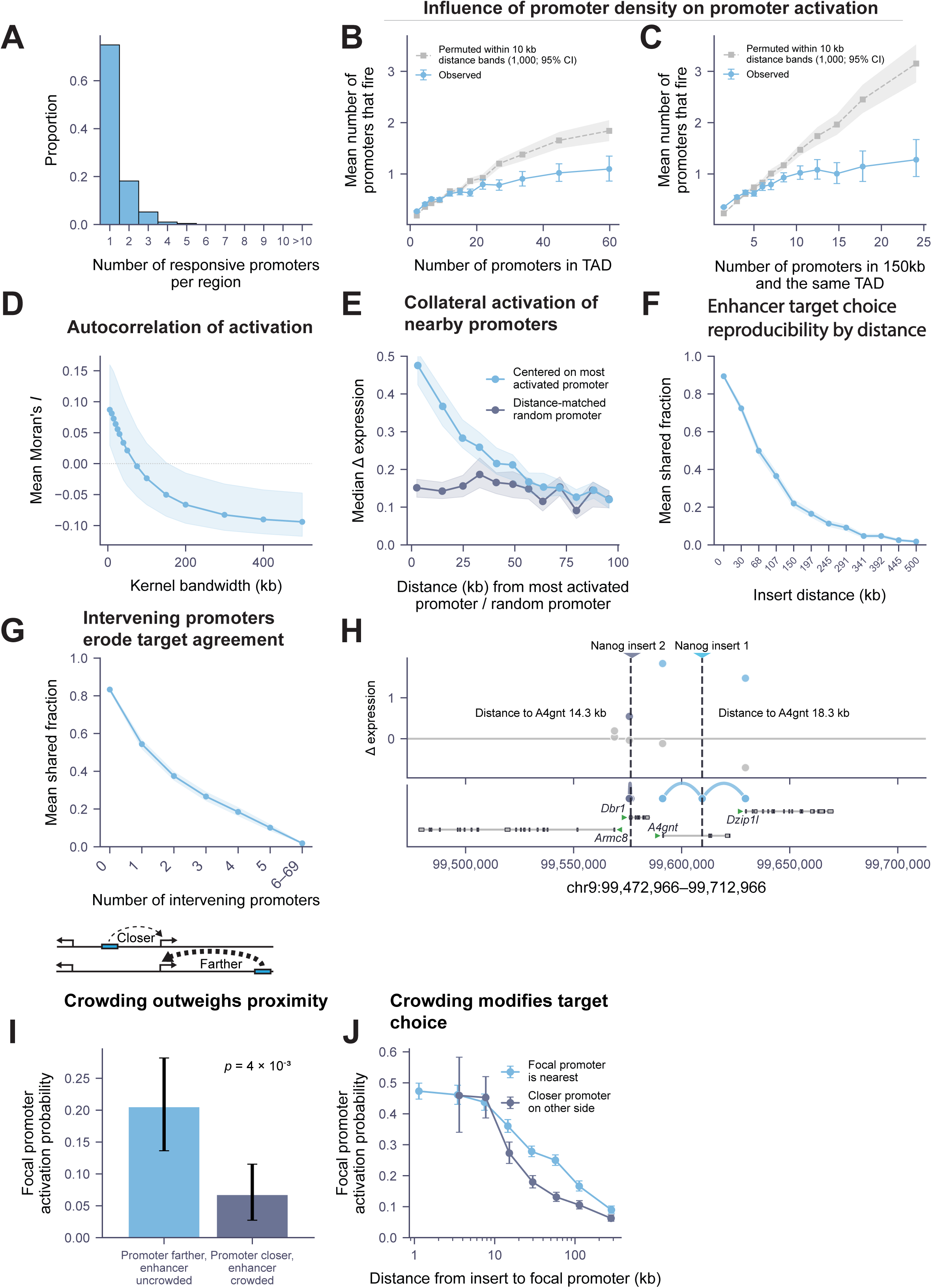
Competition among promoters shapes enhancer target choice within the local genomic landscape. **(A)** Distribution of the number of promoters activated per insertion. Number of significantly upregulated promoters per productive insertion (n = 6,724 insertions activating ≥1 promoter within 500 kb). Maximum observed, 9. **(B)** Activation count saturates as local promoter density increases. Mean promoters activated per insertion versus number of annotated promoters in the insertion’s TAD (n = 16,383 insertions). Blue, observed (binned means ± 95% bootstrap CIs). Grey, distance-matched null: promoters grouped into 10 kb distance bands and fired/not-fired labels shuffled within each band (1,000 permutations); dashed line, null mean; shading, 2.5th–97.5th percentile. Permutation tests on per-insertion slope (p ≤ 0.002) and sum of squared deviations from the permutation null (p ≤ 0.001, 1,000 permutations). **(C)** As in (B), versus promoters within 150 kb in the same TAD. Permutation tests on per-insertion slope (p ≤ 0.002) and sum of squared deviations from the permutation null (p ≤ 0.001, 1,000 permutations; n = 12,632 insertions). **(D)** Nearby promoters tend to co-activate while distant ones compete. Spatial autocorrelation (Moran’s I) of promoter activation within 500 kb, with exponentially decaying weights (w = exp(−d/λ)). x-axis, window sizes λ. Line, mean across regions; shading, 30th– 70th percentile (n = 5,501 regions). **(E)** Activation spreads locally from the strongest-responding promoter. Median Δ expression (shrunk log fold change) versus distance from the most strongly activated promoter (blue) or from a distance-matched randomly chosen promoter (grey). The center promoter is excluded from both curves. Shading, 95% bootstrap CI of the median (n = 1,879 insertions with ≥10 promoters within 500 kb). **(F)** Target choice is largely determined by genomic context. Fraction of activated promoters shared between pairs of insertions in different clonal lines (regardless of enhancer identity), binned by inter-insertion distance. Line, mean; shading, 95% bootstrapped confidence intervals (n = 13,683 pairs). **(G)** Intervening promoters reduce target concordance. As in (F), binned by the number of annotated promoters between the two insertion sites. **(H)** Example of promoter crowding at chr9:99,472,966–99,712,966. Two Nanog insertions in independent lines (dashed lines, colored by insertion). Top, Δ expression (shrunk log fold change) per promoter; colored, significant; grey, not significant. Bottom, gene models with arcs connecting significant pairs. **(I)** A paired comparison showing that crowding reduces activation even when the enhancer is closer. Each focal promoter is observed at two independent insertion sites in different clonal lines: one where it is the nearest promoter to the enhancer (uncontested, greater enhancer–promoter distance) and one where ≥2 other promoters lie closer to the enhancer (crowded, shorter enhancer–promoter distance); both insertions within the same TAD, crowded distance ≤1.5× uncontested distance. Activation probability 0.205 (95% CI 0.136– 0.282) uncontested versus 0.067 (0.027–0.115) crowded; n = 110 promoters. p = 4 × 10⁻³, two-sided exact binomial sign test. **(J)** Competing promoters reduce the effective range of enhancer activation. Activation probability versus insert–promoter distance for focal promoters with no closer promoter on the other side of the insertion (blue) versus those where a closer promoter exists on the opposite side (purple). n = 15,069 insertions, 23,751 promoters. Error bars, 95% Wilson score intervals.

The distinction between housekeeping and developmental genes is frequently invoked as a determinant of whether a promoter can respond to an enhancer, so we hypothesized that a promoter’s expression dynamics across cell types might carry information about its responsiveness. Indeed, recent work suggests that housekeeping promoters are less responsive to enhancers, operate near saturation of transcriptional output, and only reveal their enhancer dependency under conditions associated with fluctuations of core promoter regulating TFs (*8–10*, *16*). Given that the housekeeping genes are characterized by low expression variance across cell types, we turned to FANTOM5 consortium data, which allowed us to derive quantitative descriptors of promoter behavior across a broad panel of cell types. Chief among these is promoter dynamic range, which we define as the difference in log expression between a promoter’s 5th- and 95th-percentile expression states. Promoters with high dynamic range undergo larger expression shifts during differentiation than those with low dynamic range, which we reasoned might render them more susceptible to enhancer influence. Dynamic range alone does not separate housekeeping from developmental promoters, but combined with other FANTOM5-derived traits—including the Gini coefficient of promoter expression (a measure of expression inequality across cell types;(*14*, *15*))—it supports a rigorous classification. Even so, we note that the separation is purely quantitative: promoters form a smooth gradient from broadly expressed across tissues to strongly cell type-restricted, with no natural break between the two classes (Sup. Fig. 5A-F). We therefore used these continuous descriptors rather than a binary assignment to analyze the distribution of dynamic range and Gini coefficient values across the three classes of promoters.

**Figure 5.**
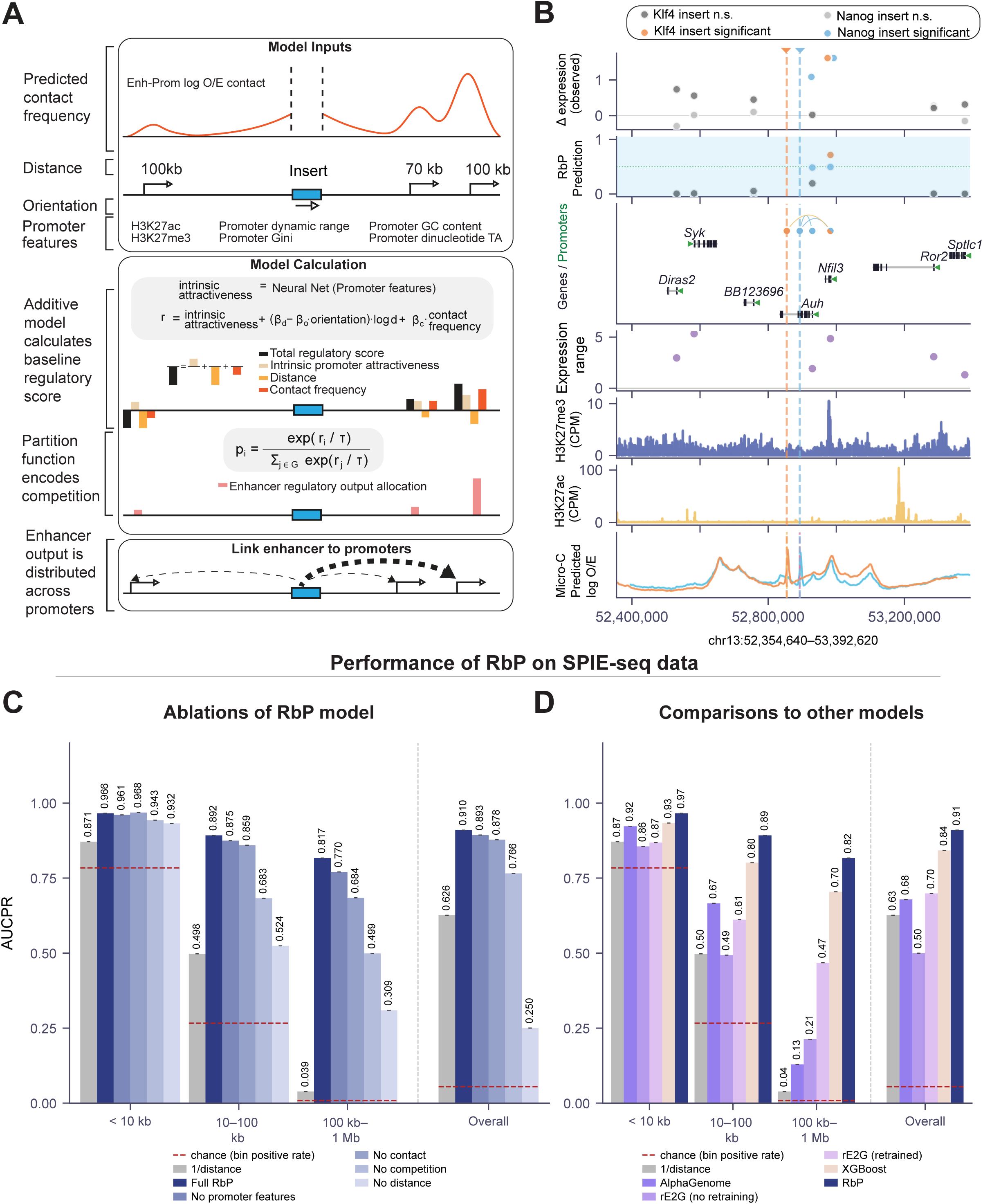
A Regulation-by-Partition model predicts enhancer target choice and outperforms existing enhancer-gene models. **(A)** RbP model schematic. Each promoter within 1 Mb is scored using four inputs: predicted enhancer–promoter contact (log observed/expected Micro-C, predicted from sequence), genomic distance, orientation of the inserted enhancer, and promoter features (H3K27ac, H3K27me3, expression dynamic range, Gini coefficient, GC content, TA dinucleotide content). A three-layer neural network maps promoter features to an intrinsic attractiveness term (no access to distance or contact). The raw score is r_i = attractiveness_i + (β_d − β_o·orientation_i)·log d_i + β_c·contact_i. Scores are passed through a partition function with temperature τ, partitioning enhancer output across candidates. Trained on chromosomes 5–19 and chrX; hyperparameters tuned on chromosomes 3–4. **(B)** RbP predictions at the *Auh–Nfil3* locus (chr13:52,354,640–53,392,620). A Klf4 (orange) and Nanog (blue) enhancer inserted in separate lines, both within Auh introns. Tracks, top to bottom: observed Δ expression (shrunk log fold change; colored, significant; grey, not significant); RbP-predicted shares (chr13 held out retrained model); gene models with arcs for significant pairs; promoter expression dynamic range; H3K27me3 and H3K27ac ChIP-seq (CPM); predicted Micro-C contact (log O/E) from each insertion site, colored by insertion. **(C)** Contribution of each RbP component. AUCPR for classifying pairs as significantly activated on held-out chromosomes 1–2, restricted to loci with ≥1 significant promoter. Split by distance (<10 kb, 10–100 kb, 100 kb–1 Mb) and pooled. Ablated models retrained without, as indicated: all promoter features, predicted contact, competition (partition function replaced by independent sigmoid), or distance. Grey, 1/distance baseline. Red dashed lines, chance AUCPR. **(D)** Comparison with existing methods on the same held-out pairs and distance bins: 1/distance, AlphaGenome predicted enhancer effect, ENCODE-rE2G (with and without SPIE-seq retraining), XGBoost trained on SPIE-seq (422 features), and RbP.

We found that the dynamic range and Gini coefficient are correlates of promoter responsiveness, with regulated promoters showing greater dynamic range and higher expression unevenness than either skipped or non-regulated promoters; the effect is particularly apparent among very distal regulated promoters (Fig. 3D,E).

We next analyzed chromatin feature enrichment of the three promoter classes across the same distance bins. Interestingly, we observed that with increasing E-P distances some features became more distinguishing between the three promoter classes. Specifically, at greater E-P distances regulated promoters showed lower H3K27ac and higher H3K27me3 compared to those that are skipped or non-regulated, while feature distributions remained broadly overlapping at short range (Fig. 3F,G). H3K9me3 showed modest enrichment at regulated long-range promoters, indicating that skipping is unlikely to be explained by promoter heterochromatinization that renders it enhancer unresponsive (Fig. 3H). CTCF and Rad21 showed enrichment at the medium- and long-range responsive promoters; responsive promoters also showed higher GC content (Fig. 3I-K).

Altogether, we conclude that among all examined promoter features, those related to measures of their developmental regulation, including dynamic range, Gini coefficient and H3K27me3 enrichment – a hallmark of repressed or poised developmental promoters – are most strongly associated with responsiveness of these promoters to enhancer activation, but that none of these features can fully explain responsiveness or skipping. Furthermore, the more apparent enrichment of features associated with promoter responsiveness at greater E-P distances can be explained by a tradeoff between genomic distance and intrinsic ‘attractiveness’ of the promoter to the enhancer: a promoter far from the insertion can only respond detectably if its intrinsic attractiveness to the enhancer compensates for the long distance penalty.

### Enhancer–promoter specificity is shaped by promoter competition within the local genomic context

Having characterized intrinsic promoter features associated with enhancer responsiveness, we next asked whether the broader genomic context—specifically the number and arrangement of nearby promoters—influences enhancer target choice. We first explored how frequently the inserted enhancer significantly upregulates expression from more than one promoter. Restricting this analysis to productive insertions—those activating at least one promoter within 500 kb—we asked how many of the available candidate promoters were activated at each site. Although these insertions had a median of 9 candidate promoters within 500 kb, the enhancer activated a single promoter in approximately 75% of cases and more than one in the remaining 25%, for a mean of 1.35 promoters activated per insertion (Fig. 4A). Target choice is therefore selective rather than a consequence of promoter sparsity: at most loci the enhancer picks one promoter from among many within reach.

How does the density of surrounding promoters affect this behavior? To address this, we returned to the full set of enhancer insertions, including nonproductive ones. Comparing the number of promoters within the enhancer’s TAD against the number that responded revealed clear diminishing returns relative to a distance-matched permutation null (Fig. 4B; permutation p ≤ 0.002 for the slope). Even in TADs containing tens of promoters, the enhancer activates only one promoter on average (Fig. 4B). A similar relationship holds when promoter density is measured within 150 kb in the same TAD, a window well inside the enhancer’s effective range, where observed activation counts again fall below the distance-matched null (Fig. 4C). Because promoters differ in their intrinsic responsiveness which correlates with developmental features (Fig. 3), this sublinear relationship could in principle reflect the supply of eligible targets rather than competition between them. We therefore repeated the analysis counting only promoters of equivalent class (i.e. developmental vs housekeeping, as approximated by the Gini coefficient), so that the available targets are matched for eligibility. Growth remained sublinear, and the per-promoter firing rate relative to its distance-based expectation declined steadily as the window filled (Sup. Fig. 6A and B). Moreover, even when we conditioned the data on regions where at least one promoter responded, the number of activated promoters plateaued at 2, regardless of how many were present within an enhancer’s reach (Sup. Fig. 6C). This pattern is consistent with a competition model in which adding promoters increases the total number of targets far less than proportionally, with one or two promoters capturing most of the enhancer output at the expense of the others.

**Figure 6.**
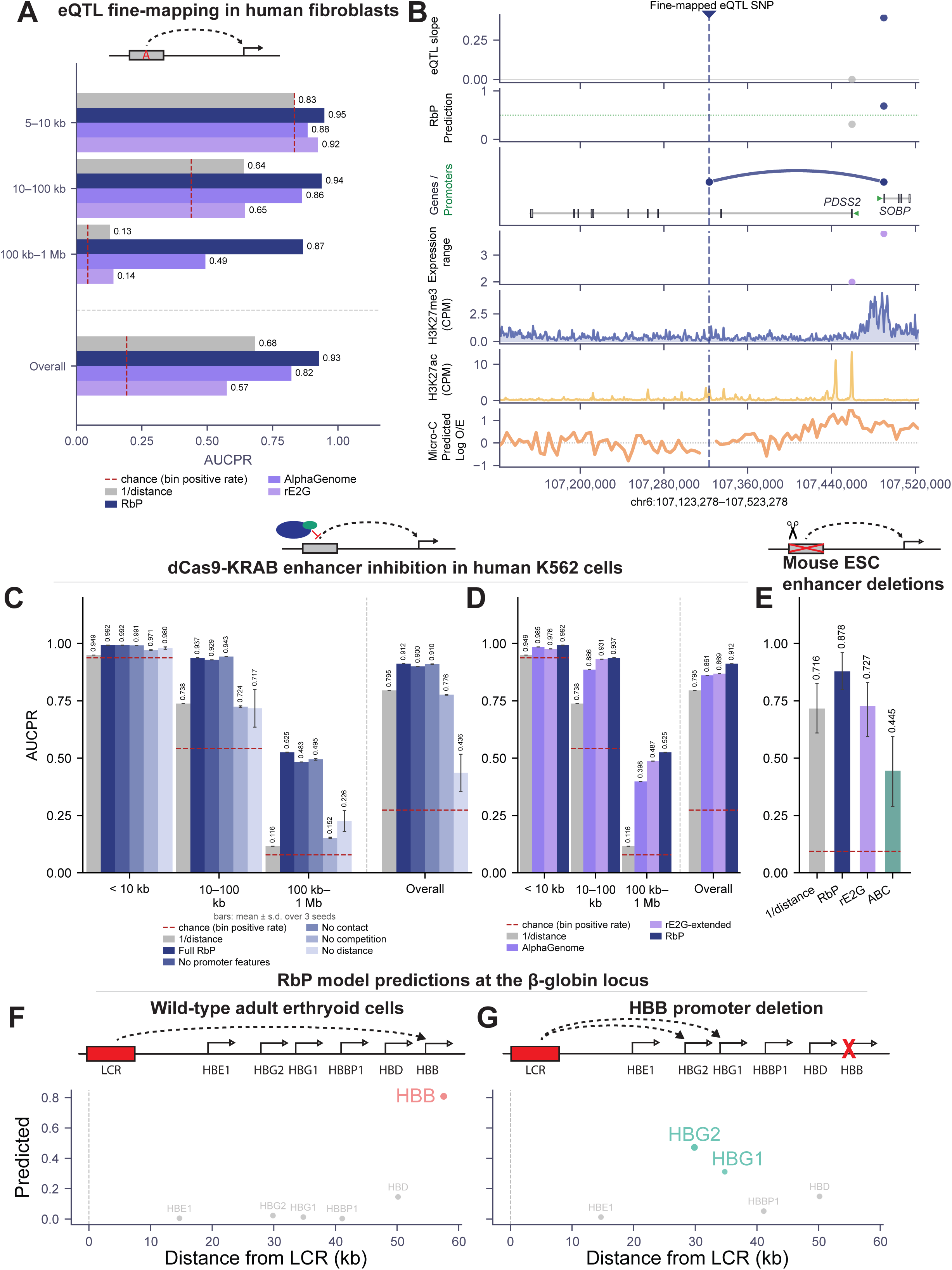
RbP generalizes zero-shot across species, cell types, and perturbation modalities. **(A)** Enhancer–gene assignment for fine-mapped eQTLs in GTEx human cultured fibroblasts. SNPs in the 95% credible set intersecting ENCODE candidate cis-regulatory elements were paired with all promoters within 1 Mb. Bars, AUCPR by SNP–TSS distance strata and overall for 1/distance, AlphaGenome, ENCODE-rE2G, and RbP (zero-shot, weights fixed from SPIE-seq). Red dashed lines, chance AUCPR. **(B)** Example locus (chr6:107,123,278–107,523,278, hg38): a fine-mapped SNP in a *PDSS2* intron whose eQTL target is *SOBP*. Tracks: eQTL effect size per gene; RbP prediction (dotted line, 0.5 threshold); gene models with the top predicted link as an arc; promoter expression range; H3K27me3 and H3K27ac (CPM) in cultured fibroblasts; predicted Micro-C contact from the SNP. **(C)** Component ablation on CRISPRi enhancer-silencing data in K562 cells (*19*, *21*, *22*, *52*). Restricted to elements with ≥1 responsive promoter. Bars, AUCPR by element–TSS distance and overall for the full model and variants omitting promoter features, contact, competition, or distance. Mean ± s.d. over 3 training seeds. **(D)** Comparison of RbP with 1/distance, AlphaGenome, and ENCODE-rE2G (extended) on the same CRISPRi benchmark. rE2G was trained on this dataset; RbP and AlphaGenome are zero-shot. **(E)** Performance on 18 validated enhancer deletions in mouse ESCs (*25–29*, *57*), scored by 1/distance, RbP, ENCODE-rE2G, and ABC. Error bars, 95% bootstrap confidence intervals over regions (4000 samples). **(F)** RbP-predicted share of LCR output at the wild-type β-globin locus in HUDEP-2 cells. Top, locus schematic; bottom, predicted share per promoter versus distance from the LCR. **(G)** As in (F), after in silico deletion of the HBB promoter.

To explore the competition further, we performed spatial autocorrelation analysis of activation effects (Moran’s I, computed across a range of window sizes). It revealed co-activation of promoters located within short distances from each other that transitions to a competitive relationship at longer distances (Fig. 4D). Consistent with this, in ∼70% of insertions activating two promoters the two targets were immediate neighbors (median zero intervening promoters, median separation 15 kb). The short-range co-activation noted above is also visible in our data directly: we noticed that when an enhancer activates a primary target, neighboring promoters often show a modest boost in expression. One such example is the *Hoxd* locus in which an inserted Klf4 enhancer skips over *Lnpk* and regulates *Evx2*, also activating *Hoxd13* (Sup. Fig. 6D). To formalize this observation, we defined two sets of promoters — the most strongly activated promoter in each region, and a distance-matched set of randomly selected promoters — and asked how the expression of their neighbors varied with P-P distance. At the shortest separations, neighbors of an activated promoter reached a median shrunk log fold change of ∼0.48, compared with ∼0.15 for distance-matched neighbors, an excess of ∼0.33 (≈1.4-fold), and this excess decayed sharply with distance, converging on the matched set by ∼60 kb (Fig. 4E). This decay tracked distance from the activated promoter more closely than distance from the enhancer, which is more consistent with a local collateral effect of activation at the primary target than with the enhancer distributing its output across several nearby promoters, though the two distances are correlated and we cannot fully separate them here. We also cannot exclude that insertions landing in permissive chromatin yield a locally stronger enhancer that raises all nearby promoters. Short-range co-activation and long-range competition need not be in conflict: competition may act on enhancer output while the local boost arises downstream of activation at the primary target.

To further probe the determinants of target specificity, we compared insertions from different clonal lines that landed near one another within the same genomic region, across inter-insert distances from 100 bp to 500 kb. Insertions separated by less than 50 kb shared the majority of their target genes (mean shared fraction ≈0.75), indicating that target choice is largely robust to small positional shifts (Fig. 4F). Beyond this separation, concordance declined gradually rather than abruptly with distance.

We asked whether this decay reflects the increasing likelihood that a promoter lies between the two insertion sites and could redirect enhancer activity. Stratifying the analysis by the number of intervening promoters, each additional promoter lowered concordance at matched distances: one intervening promoter reduced the mean shared fraction by roughly 0.1–0.2, and three or more reduced it below 0.25 across all distance bins (Fig. 4G, Sup. Fig. 6E). The decay with distance nonetheless persisted within every intervening-promoter class, indicating that intervening promoters contribute to, but do not fully account for, the distance dependence of target choice.

We next asked whether the local promoter landscape, rather than intrinsic promoter features alone, influences which promoter an enhancer activates. To test this with intrinsic attractiveness held constant, we identified promoters that received nearby enhancer insertions at two independent sites in different clonal lines: one at which the promoter was uncontested (no other promoter lay closer to the enhancer), and one at which at least two promoters lay closer (example in Fig. 4H). Because the comparison is made within the same promoter, intrinsic attractiveness is held constant by construction. We further required that the enhancer was positioned closer to the promoter in question in the crowded scenario, compared to the uncontested one, and that neither E-P pair is separated by a TAD boundary. Despite being further from the enhancer, the same promoter responded in 20.5% of cases when uncontested versus only 6.7% when it and competitors lay closer (n = 110 promoters, p = 4 × 10⁻³; Fig. 4H–4I). A comparable difference was obtained when the TAD restriction was relaxed, with the crowded enhancer again strictly closer (19.1% versus 5.0%, n = 231, p = 2 × 10⁻⁶; Sup. Fig. 6F), and when competitors were counted only within the promoter’s TAD (15.9% versus 7.1%, n = 131, p = 2 × 10⁻²; Sup. Fig. 6F).

To ask whether this effect extends beyond the matched-pair design, we binned all enhancer– promoter pairs by genomic distance and compared activation rates for enhancers inserted in crowded versus uncrowded landscapes. Crowded enhancers were less likely to activate a distal promoter even when no other promoter lay between the enhancer and its target, indicating that competition does not operate solely through interception by intervening promoters (Fig. 4J).

Together, these observations suggest that while intrinsic promoter features contribute to a promoter’s potential for regulation (Fig. 3), the local promoter landscape plays a substantial and unexpected role in determining an enhancer’s target choice.

### A regulation-by-partition model predicts enhancer target choice

Together, the preceding results identify three parameters that influence the responsiveness of promoters to an enhancer: (i) decay with genomic distance (Fig. 2); (ii) promoter features, including high expression dynamic range, Gini coefficient and higher H3K27me3/lower H3K27ac enrichment (Fig. 3); and (iii) competition among promoters within the landscape (Fig. 4). We set out to develop an interpretable, quantitative model that incorporates these insights to predict promoter responses to enhancers. We refer to it as the Regulation-by-Partition (RbP) model, since the partitioning of enhancer output allocation among promoters proves important to its performance.

The RbP model operates in two steps. First, each promoter within 1 Mb of the insertion receives a raw regulatory score, computed as a linear combination of four features: (1) log genomic distance between the insertion and the promoter, (2) predicted contact frequency, obtained from a custom neural network that predicts log observed-over-expected Micro-C contact from sequence, applied to the locus carrying the inserted enhancer rather than the reference genome, so that contacts reflect the modified sequence. Because contacts are expressed as observed over expected, the distance dependence of contact frequency is normalized out, and this term captures deviation from the distance expectation rather than contact frequency itself (see Methods), (3) enhancer orientation relative to the promoter, since piggyBac integration is not orientation-specific, and (4) intrinsic promoter features (i.e. expression dynamic range and Gini coefficient across cell types, H3K27ac and H3K27me3 levels, GC content, and TA dinucleotide content), which are weighted through a neural network to calculate intrinsic attractiveness of the promoter (Fig. 5A). Second, these raw total regulatory scores pass through a partition function, yielding values that sum to one across all candidate promoters in the landscape. The partition determines *which* promoters receive enhancer output; a separate output layer, taking these shares together with the promoter features, predicts *how much* expression changes at each one. All model parameters — the weights of the attractiveness network, the distance and contact coefficients, and the output layer — are learned jointly by training on SPIE-seq insertion events. We trained on chromosomes 5–19 and chrX, tuned hyperparameters on chromosomes 3–4, and reserved chromosomes 1–2 for final evaluation; no data from the evaluation chromosomes were used at any stage of model development. Intrinsic attractiveness is computed by a three-layer neural network over promoter features alone, which has no access to genomic distance or predicted contact, so that these terms enter only through the linear combination that forms the regulatory score. The same trained weights are used without modification for all transfer experiments described below. Altogether, the RbP provides quantitative allocation of enhancer regulatory output to the promoters within the locus, predicting enhancer target choice and expression effect (Fig. 5A).

We interpret the partition output not as the probability that a promoter is activated, but as the share of a limited enhancer output allocated to it, so that raising one promoter’s share necessarily lowers every other’s (Fig. 5A). This form was motivated by the saturation in Fig. 4B,C, where adding promoters to a landscape does not increase the total number of targets. Under this interpretation, a skipped promoter is one whose allocated share is too small to produce a detectable change in expression. The *Nfil3* locus illustrates how competition and distance interact at a single site. This locus received two independent insertions in different clonal lines: a Nanog enhancer and, more distally, a Klf4 enhancer, both landing within introns of *Auh*. The proximal Nanog insertion activated both *Auh* and the more distant *Nfil3*, whereas the distal Klf4 insertion skipped *Auh* entirely and activated *Nfil3* alone (Fig. 5B). Because enhancer identity seemingly contributes little to target choice relative to genomic position (Fig. 1G), we asked whether RbP could account for this difference from position alone. We retrained the model with chromosome 13 held out. In the model, *Nfil3* carries the higher intrinsic score of the two promoters and is predicted to form stronger contacts with an inserted enhancer, while *Auh* is favored only by proximity. At the Nanog insertion, *Auh*’s distance advantage is large enough to offset this gap, and output is divided between the two promoters (*Nfil3* 0.49, *Auh* 0.48), matching the observed dual activation. At the distal Klf4 insertion, both promoters sit far enough away that their distance penalties are nearly equal— the power-law decay compresses the difference—so the intrinsic gap dominates and output concentrates on *Nfil3* (p=0.71 versus 0.19 for *Auh*), leaving *Auh*’s share below the level at which we detect a significant change in expression.

We evaluated the RbP model on held-out chromosomes, restricting the evaluation to regions containing at least one significantly changing promoter. We report area under the precision– recall curve (AUCPR) rather than AUROC because the data are highly imbalanced — only 3– 8% of promoters near an insertion are regulated — making precision–recall the more informative metric. The full RbP model achieved an overall AUCPR of 0.91, with performance declining at greater enhancer–promoter distances (0.97, 0.89 and 0.82 for <10 kb, 10–100 kb and 100 kb–1 Mb, respectively; Fig. 5C). Since we have used a custom neural network model for contact prediction, we have further evaluated RbP performance by substituting these predictions with either experimental Micro-C measurements or contact predictions from the established AlphaGenome model (*23*, *39*). RbP performed similarly across all three contact inputs, though the two in silico contact prediction methods showed improvement over experimental contact measurements, likely reflecting lower noise in predicted than in measured contacts (Sup. Fig. 7A).

To determine which components of the RbP model carry the predictive signal, we performed ablations — removing individual model components, retraining, and reassessing performance. Partition proved essential: removing the partition function reduced overall AUCPR from 0.91 to 0.77, with the collapse most severe at long range (0.817 → 0.464 beyond 100 kb;Fig. 5C). Removing distance itself was the most damaging single ablation overall (AUCPR 0.26), confirming genomic distance as the dominant predictor. Ablating predicted contact frequency had a smaller but distance-dependent effect, reducing overall AUCPR to 0.87; the loss was confined to the mid- and long-range bins (0.89 → 0.86 at 10–100 kb and 0.817 → 0.644 beyond 100 kb), with essentially no change below 10 kb (0.97 → 0.97). The distance and contact terms are only weakly correlated in our data (Pearson r = 0.04), consistent with the observed-over-expected normalization. Removing both together reduced AUCPR to 0.13 — below either single ablation.

We next asked whether promoter features carry predictive signal. Removing all intrinsic promoter features — leaving only distance, predicted contact and orientation — reduced overall AUCPR from 0.91 to 0.89, with the loss concentrated beyond 100 kb (0.82 → 0.743). The penalty for removing promoter features grows with enhancer–promoter distance, mirroring the pattern in Fig. 3C,D,I–K: promoter identity matters most at long range, where distance alone no longer distinguishes between candidate promoters.

We then compared RbP to existing enhancer–gene prediction methods evaluated on the same held-out data. RbP (overall AUCPR 0.91) outperformed an XGBoost classifier trained on the same data on 422 genomic, epigenomic, expression and contact features (0.85), despite using far fewer parameters and an interpretable architecture. RbP also outperformed AlphaGenome (0.68), the rE2G model applied zero-shot (0.50) and an rE2G retrained on SPIE-seq data (0.70) (Fig. 5D). The advantage of RbP was most pronounced beyond 100 kb (0.82 versus 0.73 for XGBoost, 0.13 for AlphaGenome, and 0.47 for retrained rE2G), the regime in which target choice is least determined by proximity alone and where partition-based scoring is most informative. Because the evaluation above is restricted to loci containing at least one significantly changing promoter, models that rarely predict regulation could appear penalized. To control for this selection effect, we re-evaluated all models conditioned on at least one promoter being called as regulated, ensuring each method’s predictions are compared only where it also predicts activity. RbP retained a clear advantage (AUCPR 0.915 versus 0.748 for AlphaGenome, 0.716 for retrained rE2G, 0.558 for zero-shot rE2G, and 0.769 for distance alone; Sup. Fig. 8).

### A Regulation-by-Partition model generalizes across species, cell types and perturbations

To test whether the principles captured by RbP generalize beyond our assay, we applied the model zero-shot — without retraining — to four independent datasets spanning different species, cell types, and perturbation modalities.

We first asked whether RbP could predict the regulatory targets of natural human genetic variation. We compiled fine-mapped eQTL SNPs from GTEx cultured fibroblasts, retaining only variants that intersect ENCODE-annotated cis-regulatory elements, and scored all promoters within 1 Mb of each SNP. RbP achieved an overall AUCPR of 0.93, outperforming both AlphaGenome (0.82) and the published ENCODE-rE2G model (0.58), with the advantage most pronounced beyond 100 kb (0.78 versus 0.49 for AlphaGenome and 0.14 for rE2G; Fig. 6A). This advantage holds after correcting for locus ascertainment (Sup. Fig. 9A). An illustrative case occurs at the *PDSS2/SOBP* locus, where a fine-mapped SNP falls within an intron of *PDSS2*, yet its eQTL target is the more distal *SOBP* — a clear instance of enhancer skipping over the host gene. RbP correctly identifies *SOBP* as the top-ranked target (Fig. 6B), consistent with competition favoring a more attractive promoter despite the distance penalty.

We next applied RbP to published CRISPRi enhancer-silencing datasets in human K562 cells (*19*, *21*, *22*, *52*), using chromatin marks mapped in K562 (*53*, *54*) and intrinsic promoter characteristics estimated from FANTOM5 CAGE data in human cells (*55*, *56*); promoter sequence features were derived from the hg38 reference genome. Model weights were unchanged from training on SPIE-seq. These datasets thus represent orthogonal validation in a different cell type and species, queried with loss rather than gain of enhancer function.

RbP scores which promoter an element acts on, not whether an element is an enhancer. We therefore restricted the benchmark to perturbed regions in which at least one promoter changed significantly upon CRISPRi and evaluated all methods on the task of assigning the responsive promoter among candidates within the same window. Within this framing, the competition ablation reproduced in K562 what we observed in SPIE-seq: removing the partition reduced overall AUCPR from 0.91 to 0.81, with the collapse most severe beyond 100 kb (0.52 → 0.19), indicating that the competitive step carries the transferable signal (Fig. 6C). Predicted contact contributed comparatively little (0.912 with versus 0.910 without), as did promoter features (0.900 without). On the active-enhancer K562 benchmark, RbP (overall AUCPR 0.91) outperformed AlphaGenome (0.86), rE2G-extended which was trained on this dataset (0.87) and a 1/distance baseline (0.80; Fig. 6D). The advantage was most pronounced beyond 100 kb (0.52 versus 0.40 for AlphaGenome, 0.49 for rE2G and 0.12 for 1/distance; Fig. 6D). This lead holds after correcting for locus ascertainment (Sup. Fig. 9B) and extends to ranking accuracy (Sup. Fig. 9C). Separately, crowded enhancers in the K562 data showed reduced reach, consistent with competition limiting enhancer range across systems (Sup. Fig. 9D).

We further tested the model on 18 validated enhancer knockouts in mouse ESCs (*25–29*, *57*). Although this dataset is small, RbP performed well zero-shot (AUCPR = 0.878), outperforming rE2G (0.727), the ABC model (0.445) and a 1/distance baseline (0.716) (Fig. 6E), despite representing the opposite perturbation type — deletion rather than insertion.

As a final test, we turned to the β-globin locus, a canonical example of promoter competition for a shared enhancer. The locus control region (LCR) activates the adult β-globin gene (*HBB*) in adult erythroid cells, but deletions that remove the *HBB* promoter redirect LCR output to the fetal γ-globin genes (*HBG1/2*), a phenomenon observed both in patients with hereditary persistence of fetal hemoglobin (HPFH) and in engineered erythroid cell models (*5–7*). We asked whether RbP could recapitulate this switch. We applied the model to the wild-type locus and to a version carrying an *HBB* promoter deletion, and in each case the top-ranked target matched the experimental outcome: *HBB* dominated in the native context, while *HBG* became the primary target upon *HBB* promoter deletion (Fig. 6F,G).

Taken together, the successful transfer of the RbP model across species, cell types, perturbation modalities, and natural genetic variation suggests that distance-dependent partitioning of a fixed enhancer budget among promoters is a conserved feature of mammalian gene regulation.

## Discussion

Enhancers act across genomic windows that typically contain many candidate promoters, and the question of how one is chosen has usually been posed in the context of the enhancer–promoter pair: either the two are compatible (*16*, *17*, *58*), or an insulating structure holds them apart (*33*, *40*, *41*, *43*, *59*) or, as proposed recently, enhancer target choice is dictated by the intrinsic promoter responsiveness (*9*). Our data argue for a different model. We show that distinct enhancers inserted at the same site activate nearly the same genes (Fig. 1G), whereas insertions of the same enhancer diverge progressively in their targets as they are separated: concordance falls below half beyond ∼100 kb, and each intervening promoter lowers it further at matched distance (Fig. 4F). The number of promoters activated saturates near one, regardless of how many lie within reach (Fig. 4B,C), and the probability that the nearest promoter is the target falls as competitors accumulate (Fig. 4I). Together, these observations indicate that enhancer target choice is an emergent property of the locus rather than an inherent feature of an enhancer–promoter pair. Its quantitative logic — distance-weighted partitioning of the enhancer output among promoters of differing attractiveness — is captured by the RbP model well enough to transfer, without retraining, across species, cell type, and direction of perturbation.

This enhancer output partition reframes target specificity as a relative rather than an absolute property: whether a promoter responds depends not on its own features alone but on the features of the alternatives available to the enhancer. Consistent with this notion, feature ablations in the RbP model illustrate that removing competition affects model performance much more strongly than removing intrinsic promoter features (Fig. 5C,D). Importantly, we interpret the partition function output of RbP not as the probability that a promoter is activated, but rather, the relative share of a limited enhancer output allocated to it. As an enhancer distributes its finite regulatory output among the promoters within its reach, raising one promoter’s share necessarily lowers every other’s (Fig. 5A). This framework supplies a natural account of promoter skipping. Under a competitive model, a skipped promoter need not be intrinsically incompatible with the enhancer or shielded from it; it may simply be outcompeted by a more attractive target elsewhere in the neighborhood. In this context, skipping is not a categorical failure to respond, but an allocated share too small to produce a detectable change in expression. We thus propose that enhancer target choice is a conditional phenomenon — while it depends on intrinsic promoter responsiveness, distance and contact, the same enhancer-promoter pair can behave differently depending on which other promoters lie within range. This is illustrated well by the β-globin locus example, where RbP ranks HBB over the nearer HBE1 and HBG1/2 in the wild-type locus — the opposite of what distance predicts — and reallocates that share to HBG when the HBB promoter is deleted, matching the switch observed in HPFH and in engineered erythroid models (Fig. 6F,G; ref. *7*).

The RbP model also explains why the promoter-intrinsic features that distinguish regulated from skipped promoters emerge only at long enhancer–promoter distances (Fig. 3C,D,I–K). At short range the distance term dominates and shares are large; at long range the distance penalty compresses shares toward the detection limit, such that small differences in intrinsic attractiveness determine which promoters clear it. The same dependency appears in the RbP model feature ablations (Fig. 5C,D), where the penalty for removing promoter features grows with distance, although it remains fairly modest relative to ablations of distance itself or partition.

Our framework offers a simple account of a persistent discrepancy in the field: enhancer–promoter specificity is readily apparent at endogenous loci, yet largely absent in MPRA assays (*16*, *17*). In those assay designs the enhancer is presented with one promoter, in an episome or an otherwise empty regulatory landscape. With no competitors, the entire output is allocated to the only available target — a direct prediction of our model, and the reason such assays report promoter boostability faithfully while finding almost no specificity. A minimal assay for enhancer–promoter compatibility would need at least two promoters competing within the same landscape. By the same argument, the apparent incompatibility of skipped promoters at endogenous loci need not reflect biochemical mismatch – it is expected whenever a more attractive promoter is present. Nonetheless, we do not claim to disprove the existence of biochemical compatibility. Rather, to the extent that compatibility is visible in our data, it behaves as a property of the promoter rather than of the pair: intrinsic promoter attractiveness enters RbP as a single scalar, no enhancer-by-promoter interaction term is required, and target choice shifts with position and with the promoter landscape rather than with the enhancer identity (Fig. 1G, 4F, 4H). We note that because we tested only five enhancers in SPIE-seq, all strongly active in mESC, we cannot exclude the possibility that other types of enhancers in different cellular contexts carry genuine pairwise preferences that our panel does not sample. We further acknowledge that RbP parameters may be tuned to enhancers of specific classes and may thus not universally capture quantitative logic of long-range gene regulation across all cell types and species. Nonetheless, high performance of RbP on human eQTL data from fibroblasts and CRISPRi datasets from K562 cells, in which many different distal elements were genetically or epigenetically altered, suggest that our observations are generalizable across diverse enhancers, cell types and mammalian species. Thus, while enhancer sequence features determine when, where and how much an enhancer is active, they may be of relatively minor importance for the target choice.

RbP requires only that some quantity provided by the enhancer is limiting; it does not specify what this quantity is. Consequently, RbP remains compatible with several recently proposed promoter competition models, such as transcript-dependent insulation (*60*), which could be accommodated within RbP through the contact term rather than the promoter score: if an intervening promoter and its transcriptional unit reduce the enhancer’s contact with promoters beyond it, the enhancer’s output is redistributed toward the intervening promoter without any change to the intrinsic attractiveness of either. Our results are also not in conflict with the multi-way regulatory hub model at the globin loci, where enhancers contact several promoters without mutual exclusivity, yet these promoters compete (*61*, *62*). Further, our observation of short-range collateral boosting at promoters in vicinity to the activated target (Fig. 4E) may be understood in the context of the hub model, in which multiple nearby promoters share a local regulatory environment and the activation of the target promoter can boost its neighbors. In agreement, short-range collateral boosting decays with distance from the activated promoter rather than from the enhancer itself (Fig. 4E). Short-range synergy and long-range competition are therefore not in conflict — competition operates on enhancer output, while the local boost is a consequence of activation at the primary target.

The practical implication of our model is that the gene affected by a regulatory element is not a fixed annotation but a function of the landscape it sits in: rearrangements, deletions, and structural variants that change the set of competitors should redirect enhancer output without altering the enhancer at all, as they do at β-globin. Consistent with this, RbP applied zero-shot to fine-mapped eQTLs in human fibroblasts correctly identified distal target genes — including cases such as the *PDSS2/SOBP* locus where the causal variant skips the host gene entirely (Fig. 6A,B) — outperforming models that score enhancer–gene pairs independently. Predicting the consequences of non-coding variation may therefore require modeling the whole neighborhood rather than scoring enhancer–gene pairs one at a time.

## Acknowledgments

We thank the members of the Wysocka laboratory for critical reading of the manuscript. We also thank A. Boettiger for critical feedback on the manuscript and L.-F. Chen for sharing the original splinkerette sequencing protocol. AI assistance was used in the preparation of this work. Anthropic’s Claude was employed for text editing and proofreading. Code was drafted by hand; Claude was used to accelerate the debugging process. The authors take full responsibility for all content presented in this manuscript.

## Author contributions

Conceptualization: JV, JW, TS

Methodology: JV, JW, TS

Investigation: JV

Visualization: JV

Funding acquisition: JW

Project administration: JW

Supervision: JW

Writing – original draft: JV, JW

Writing – review C editing: JV, JW, TS

## Data, code, and materials availability

All data needed to evaluate the conclusions in this study are present in the main text, the supplementary materials, and/or the Gene Expression Omnibus (accession number pending).

## Supplementary figures

**Supplementary Figure 1.**
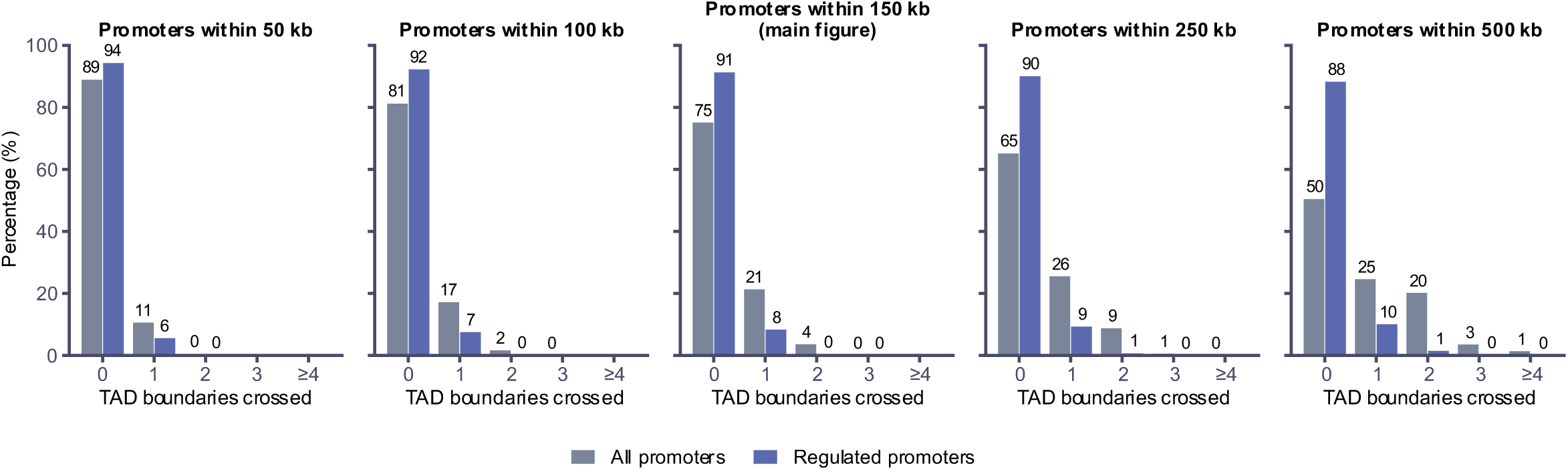
Regulatory effects and TAD boundaries. Distribution of TAD boundaries between insertion and promoter, for insertions with ≥1 significantly regulated promoter, at distance cutoffs of 50, 100, 150, 250, and 500 kb. Bars, percentage of all promoters (grey) and significantly regulated promoters (blue) at 0, 1, 2, 3, or ≥4 boundaries. The 150 kb cutoff corresponds to Fig. 2D. TAD boundaries from Bonev et al. 2017 (ref. *44*).

**Supplementary Figure 2.**
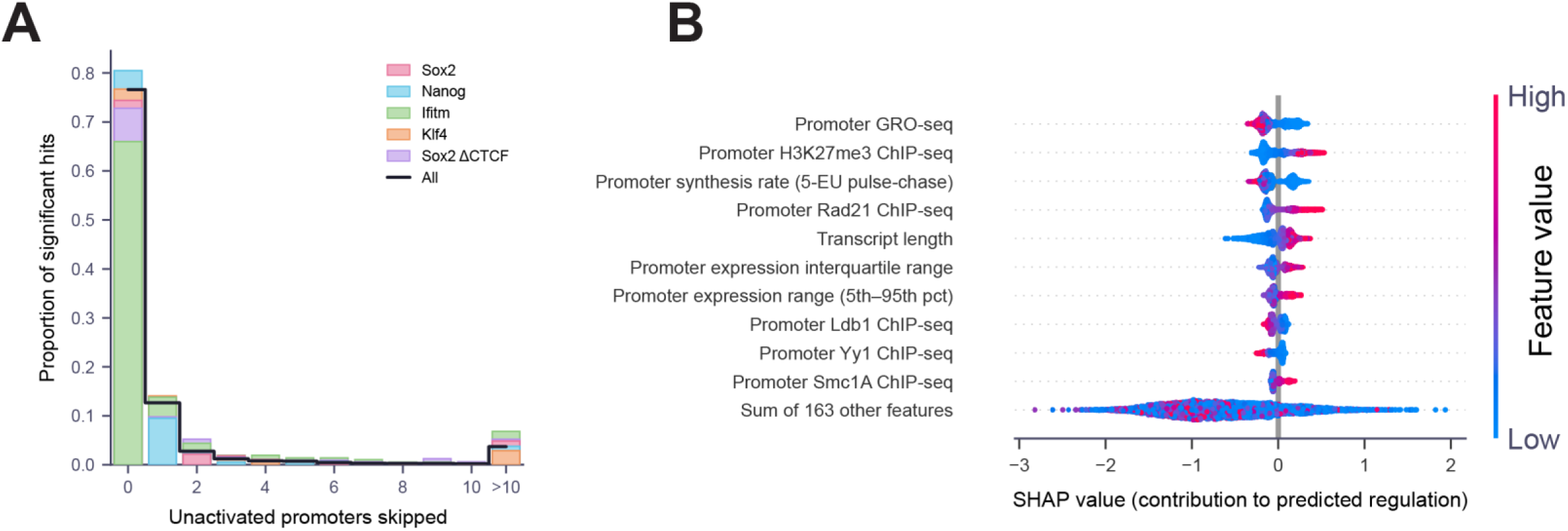
Promoter skipping and feature importance. **(A)** Number of unactivated promoters between an inserted enhancer and each promoter it significantly upregulates (0 = nearest promoter activated). Bars colored by construct; black outline, all pooled. Final bin, >10 skipped promoters. **(B)** SHAP summary for a gradient-boosted tree predicting activation from promoter features. Top ten features ranked by mean absolute SHAP value; bottom row, remaining 163 features aggregated. Each point, one promoter–insertion pair; color, feature value (blue, low; magenta, high); x-axis, SHAP contribution to predicted activation probability.

**Supplementary Figure 3.**
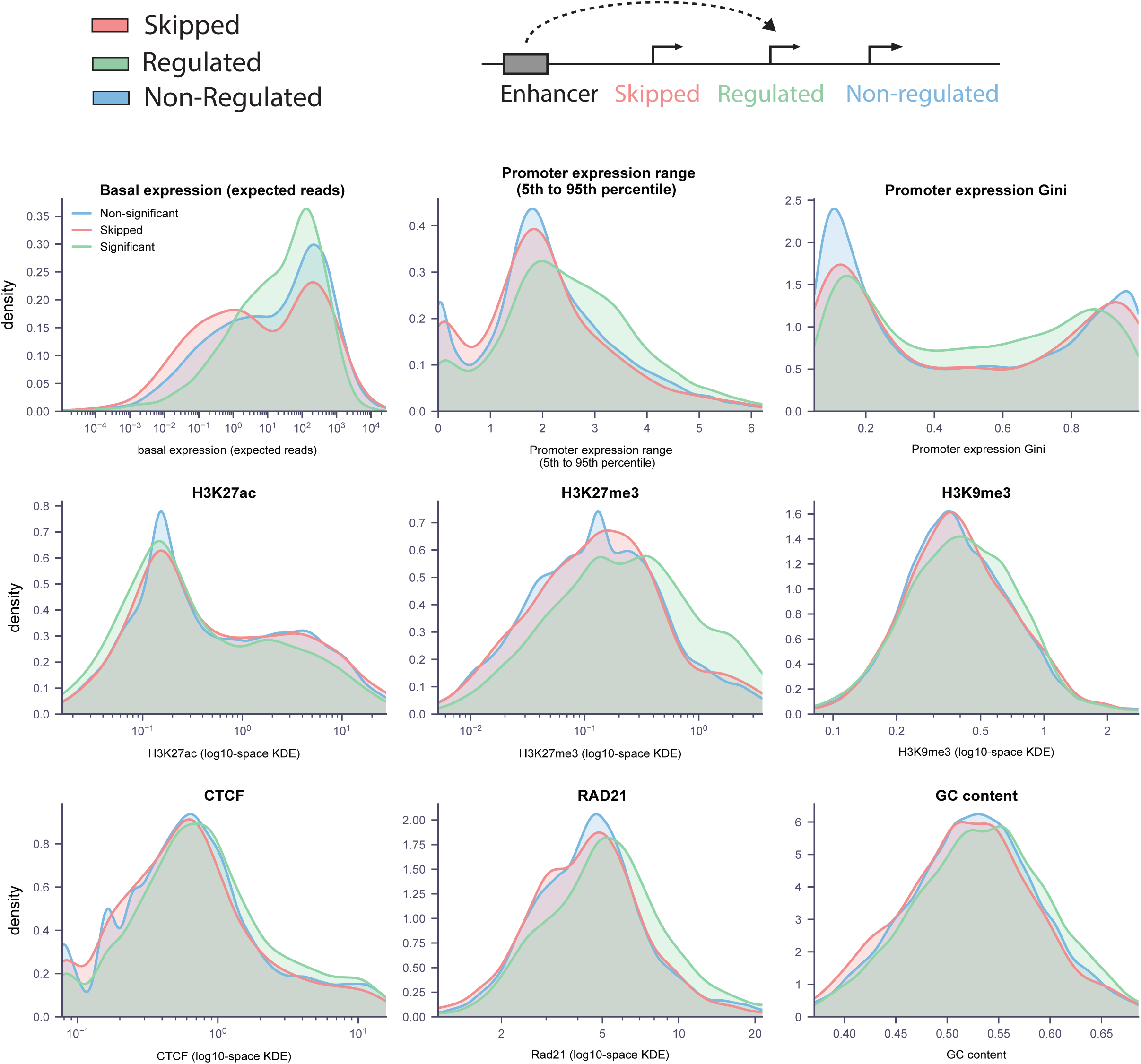
Features of regulated, skipped and non-regulated promoters. **Top:** Promoter classes as in Fig. 3. Regulated (green), significantly upregulated; skipped (pink), not upregulated but between inserted enhancer and a regulated promoter; non-regulated (blue), neither. **Bottom:** Kernel density estimates for each class. Features: H3K27ac, H3K27me3, H3K9me3, CTCF, Rad21, GC content, log basal expression, expression range (FANTOM5, p95 – p5), and expression Gini coefficient. Kolmogorov–Smirnov D and p-values shown for regulated vs skipped and regulated vs non-regulated.

**Supplementary Figure 4.**
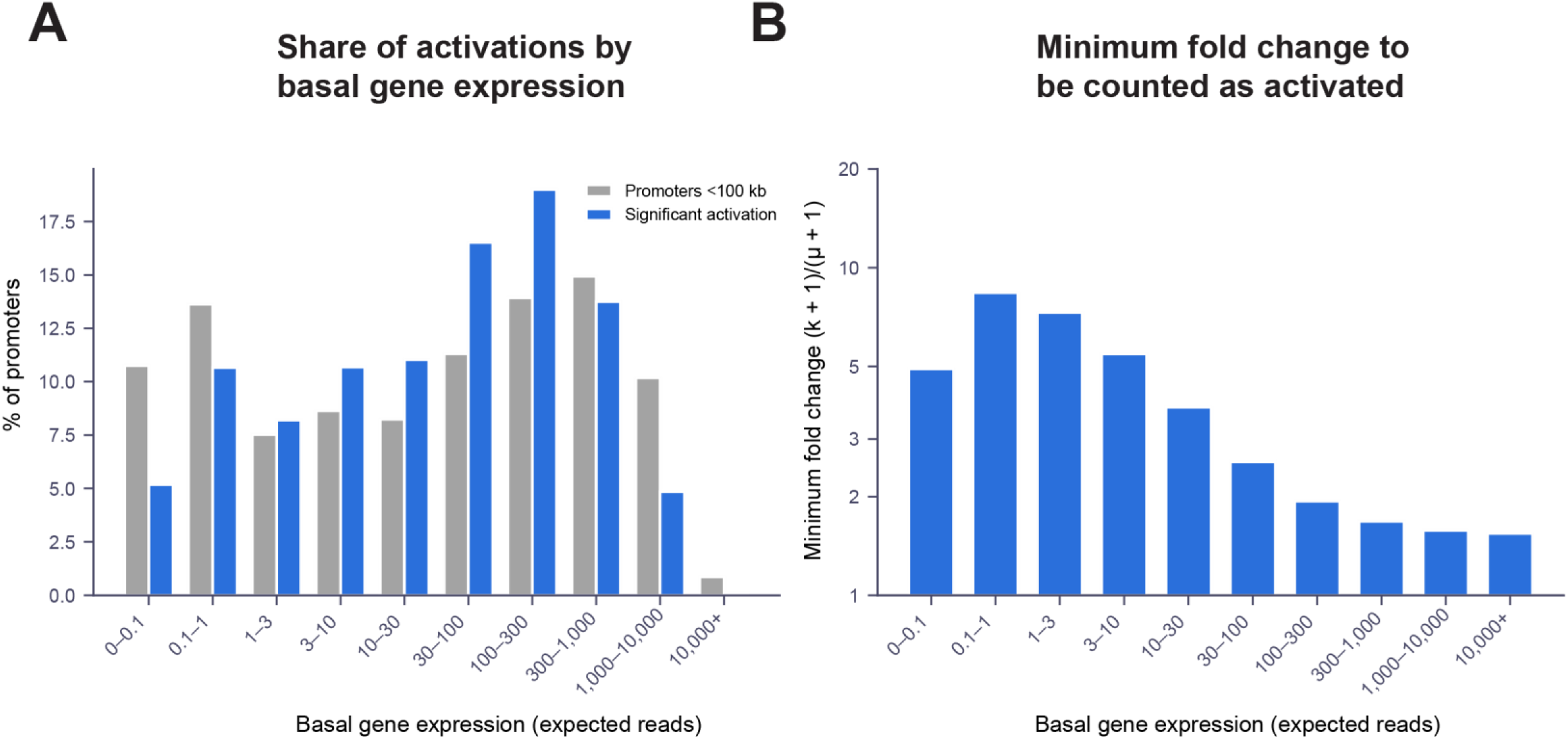
Detection sensitivity and expression of activated promoters. **(A)** Basal expression (expected read count) of significantly activated promoters (blue) versus all promoters within 100 kb of an insertion (grey). Bars, percentage per expression bin. **(B)** Minimum fold change for significant activation versus expected read count (log-binned). Fold change is (k + 1)/(μ + 1), where k is the smallest count reaching significance and μ is the expected count.

**Supplementary Figure 5.**
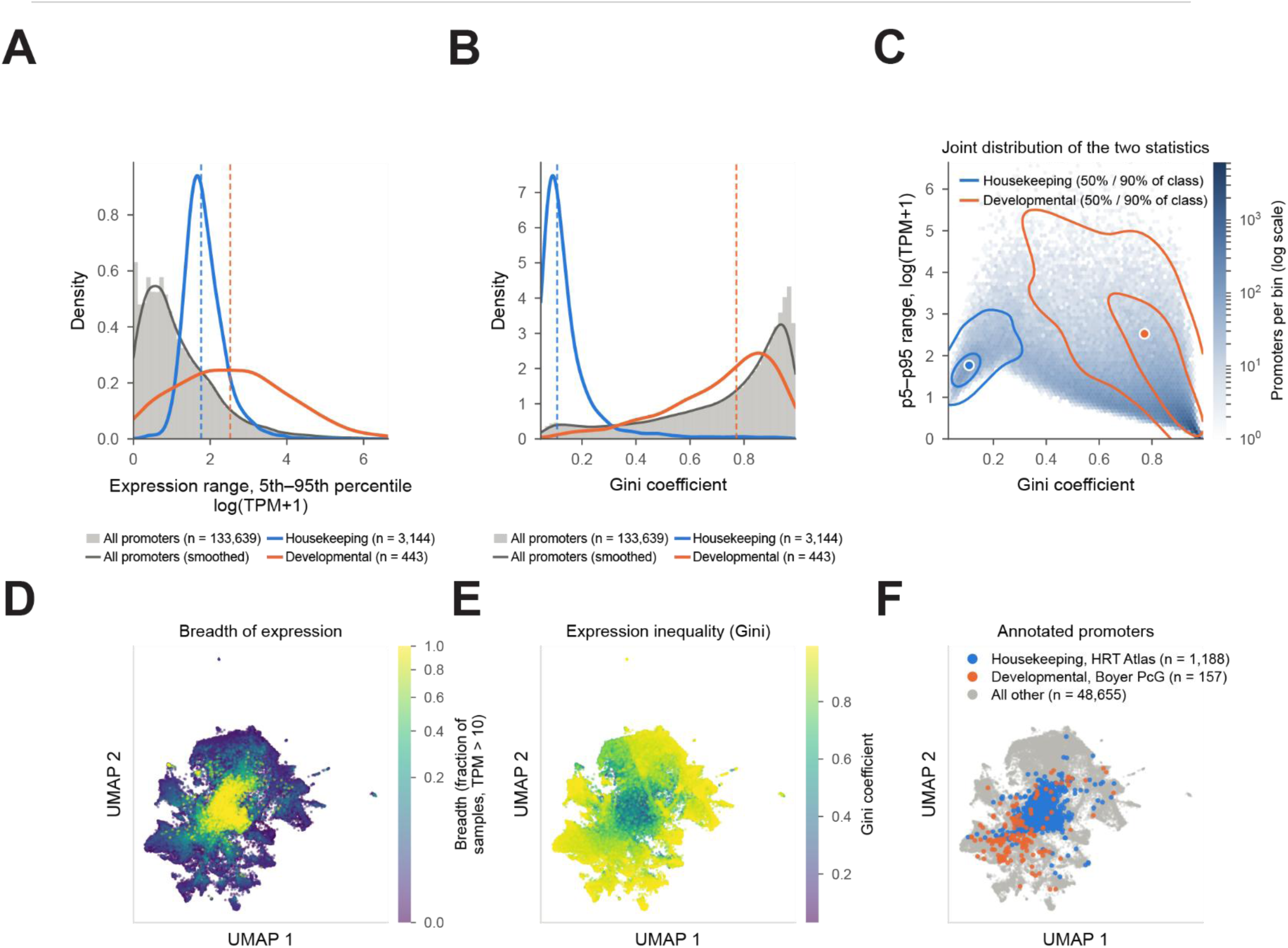
Gini coefficient versus dynamic range for classifying promoter types. **(A)** Distribution of promoter dynamic range (5^th^ to 95^th^ percentile expression log(TPM+1) across samples) for all CAGE promoters (grey, n = 133,639), housekeeping from Hounkpe et al. 2021 (blue, n = 3,144, ref. *63*) and developmental from Boyer et al. 2006 (orange, n = 443, ref. *64*) promoters. Dashed lines, class medians. **(B)** As in (A), for Gini coefficient. **(C)** Joint distribution. Hexagonal bins, all promoters (log-scale density). Contours, 50% and 90% of housekeeping (blue) and developmental (orange); white points, class medians. **(D–F)** UMAP of 50,000 CAGE promoters (cosine distance on z-scored log(TPM+1), 1,073 FANTOM5 mouse samples), colored by **(D)** expression breadth (fraction of samples with TPM > 10), **(E)** Gini coefficient, **(F)** annotation class (housekeeping, blue; developmental, orange; other, grey).

**Supplementary Figure 6.**
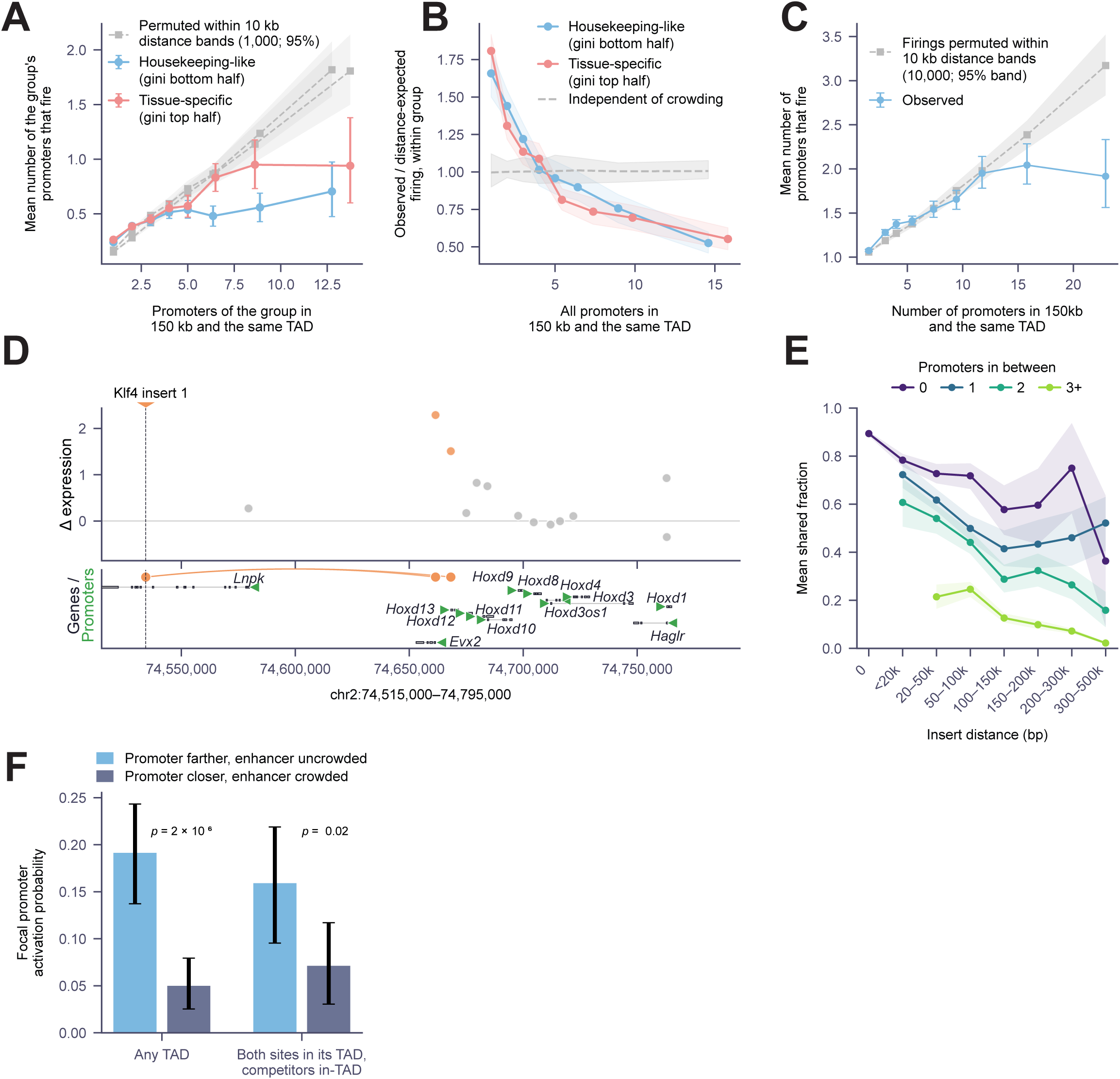
Number of activated promoters plateaus regardless of promoter class. **(A)** Mean number of same-class promoters activated versus number of same-class promoters within 150 kb and the same TAD. Promoters split by Gini coefficient: housekeeping-like (bottom half, blue) and tissue-specific (top half, red). Grey, distance-matched null (1,000 permutations shuffling labels within 10 kb bands); dashed line, null mean; shading, 95% confidence interval (CI). **(B)** Per-promoter firing rate relative to distance-based expectation, by promoter class, as a function of total promoter density (all promoters within 150 kb, same TAD). Dashed line at 1, expected if firing were independent of crowding; shading, 95% CI. **(C)** Activation count saturates as local promoter density increases. Mean promoters activated per insertion versus number of annotated promoters in the insertion’s TAD conditioned on active inserts (n = 5,130 insertions). Blue, observed (binned means ± 95% bootstrap CIs). Grey, distance-matched null: promoters grouped into 10 kb distance bands and fired/not-fired labels shuffled within each band (1,000 permutations); dashed line, null mean; shading, 2.5th–97.5th percentile. Permutation tests on per-insertion slope (p ≤ 0.002) and sum of squared deviations from the permutation null (p ≤ 0.001, 1,000 permutations). **(D)** Example of collateral activation at the *Hoxd* locus (chr2:74,515,000–74,795,000). Top, Δ expression (shrunk log fold change) per promoter after Klf4 enhancer insertion (dashed line); orange, significant; grey, not significant. Bottom, gene models with promoters (green). **(E)** Fraction of target genes shared between insertion pairs from different lines landing in the same region, binned by inter-insertion distance and stratified by number of intervening promoters (0, 1, 2, 3+). Shading, 95% CI. **(F)** Activation probability for the same promoter when the nearby insertion is uncrowded (no closer promoter; light blue) versus crowded (≥2 closer promoters; dark blue). Left, all pairs (n = 231 promoters, p = 2 × 10⁻⁶); right, both sites within the focal promoter’s TAD (n = 131, p = 0.02).

**Supplementary Figure 7.**
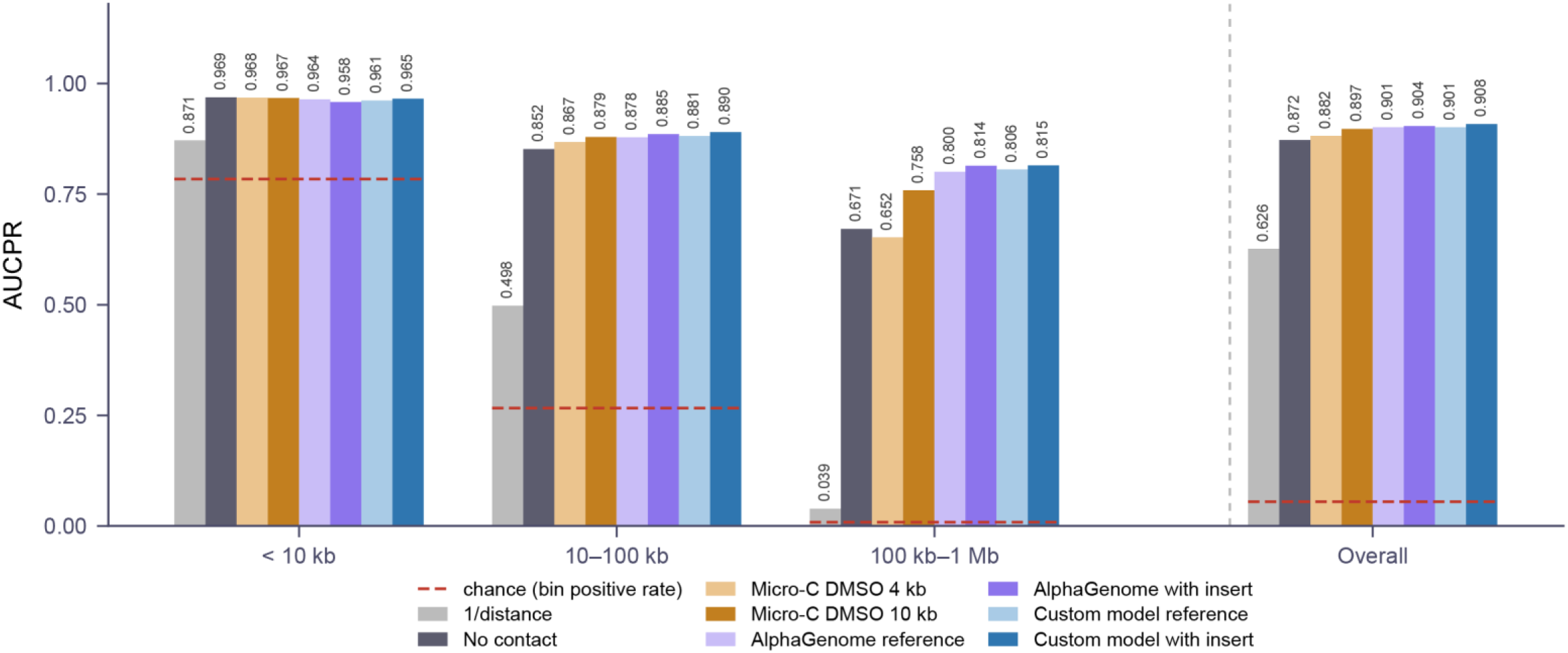
RbP performance with different contact data sources. AUCPR on held-out chromosomes 1–2, by distance stratum (<10 kb, 10–100 kb, 100 kb–1 Mb) and overall. Contact sources: experimental Micro-C (DMSO) at 4 kb or 10 kb resolution (orange); AlphaGenome predictions from reference or insert-bearing sequence (purple); Custom contact prediction model predictions from reference or insert-bearing sequence (blue).

**Supplementary Figure 8.**
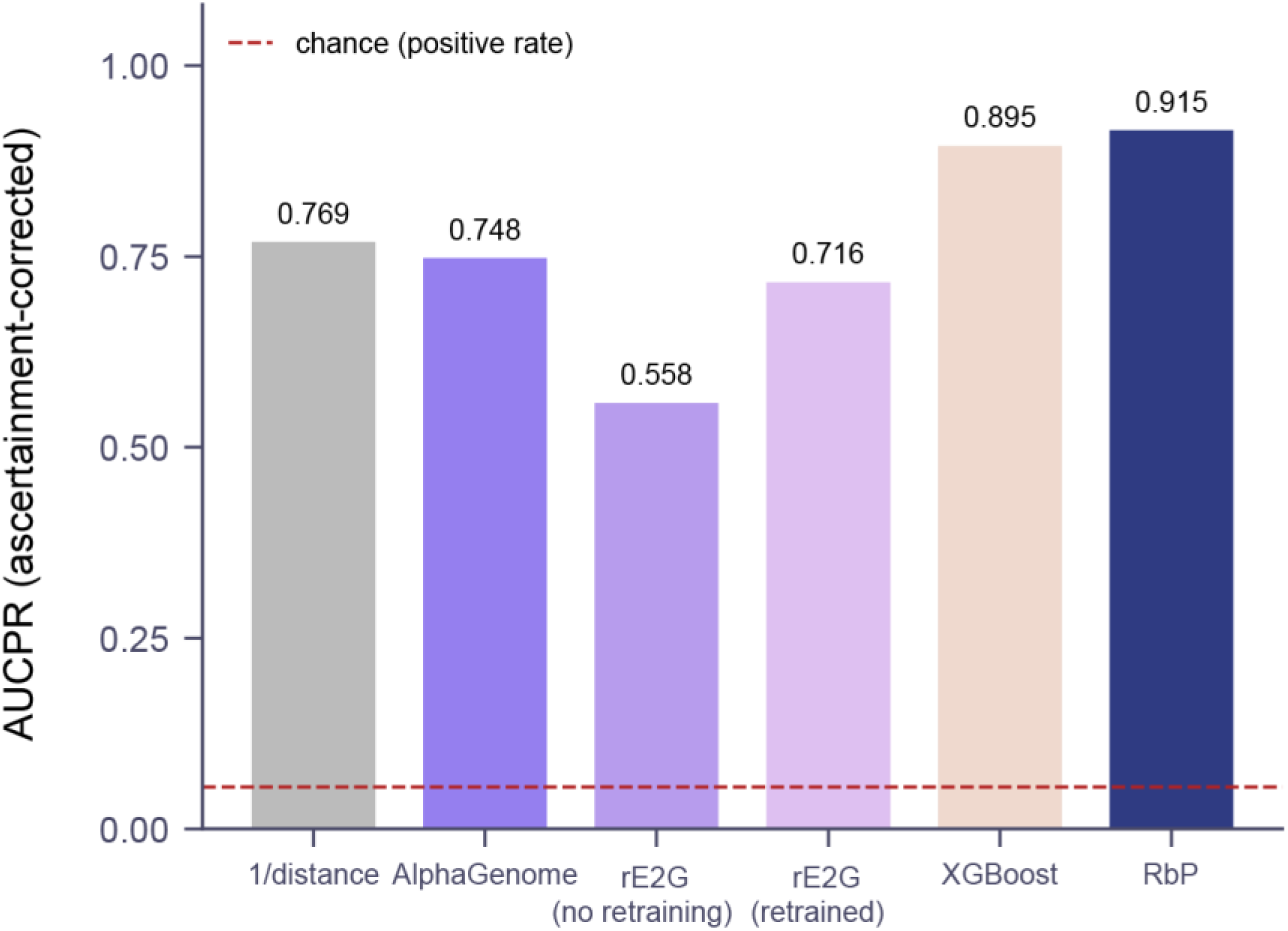
Model comparison after correcting for locus ascertainment. Evaluation loci were selected to contain at least one regulated promoter. To account for this, each model’s pair scores were conditioned on the locus containing ≥1 positive, p_i_ / (1 − ∏ⱼ(1 – pⱼ)), using noisy-OR aggregation over all candidate promoters at the locus. AUCPR on held-out chromosomes 1–2. Dashed line, positive rate.

**Supplementary Figure 9.**
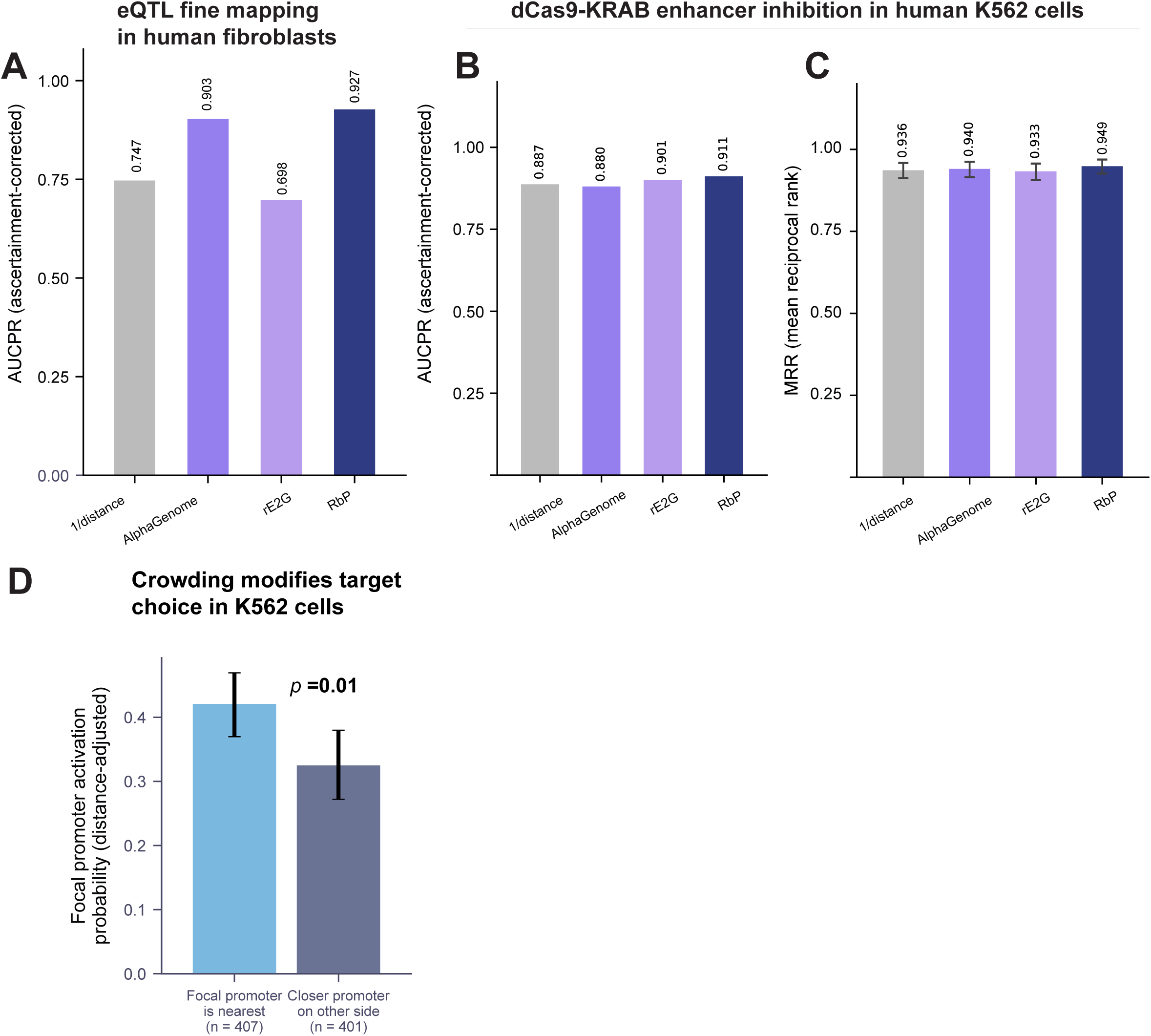
Ascertainment-corrected benchmarks, ranking accuracy, and competition in K562. **(A)** Locus-ascertainment–corrected AUCPR on fine-mapped GTEx fibroblast eQTLs. Candidate promoters reconditioned to match positive-set ascertainment (noisy-OR aggregation, p_i_ / (1 − ∏ⱼ(1 − pⱼ)). **(B)** Same reconditioning on the K562 CRISPRi benchmark. Bars as in (A). **(C)** Promoter ranking accuracy. Mean reciprocal rank (MRR) of the responsive promoter among all candidates per window on the K562 benchmark. Error bars, bootstrap 95% CI. **(D)** Distance-adjusted activation probability for focal promoters that are the nearest promoter to the element (n = 407) versus those with a competing promoter closer on the opposite side (n = 401). p = 0.01, distance-adjusted logistic regression with element-clustered standard errors.

## Materials and Methods

### Mouse ESC Culture

F121 mouse embryonic stem cells were maintained on gelatin-coated plates in N2B27 medium, consisting of a 1:1 mixture of DMEM/F12 and Neurobasal medium supplemented with N2 and Gem21 (B27 equivalent). The medium was further supplemented with two small-molecule inhibitors (2i): the MEK1/2 inhibitor PD0325901 (0.8 µM) and the GSK3β inhibitor CHIR99021 (3.3 µM), along with leukemia inhibitory factor (LIF; produced in-house and used at an empirically determined concentration sufficient to maintain pluripotency) and L-ascorbic acid (64 µg/ml).

### Enhancer selection

Four pluripotency-associated enhancers were selected for ectopic insertion: the Sox2 control region (SCR) chr3:34,756,186-34,761,866 mm10, a Nanog enhancer chr6:122,701,553-122,704,197 mm10, an enhancer at the Ifitm locus chr7:140,958,484-140,964,093 mm10, and a Klf4 enhancer chr4:55,475,057-55,478,716 mm10. Each fragment was amplified from mouse genomic DNA and cloned between the piggyBac 5’ and 3’ inverted terminal repeats of the donor plasmid, such that the enhancer fragment constituted the entire mobilizable cassette. The GFP and RFP cassettes on the donor and transposase backbones lie outside the ITRs and are therefore not mobilized, serving only to identify transfected cells during sorting. Additionally a variant of the sox2 control region with all potential CTCF sites removed was synthesized as in brosh et al 2023.

### PiggyBac Transfection and Clone Isolation

F121 cells were transfected in suspension at a density of 400,000 cells per well in a 24-well plate using Lipofectamine 2000 at a 3:1 ratio of Lipofectamine (µl) to DNA (µg). A total of 800 ng of DNA was transfected per well, consisting of piggyBac donor and transposase plasmids at a 50:1 molar ratio (donor:transposase). The donor and transposase plasmids carried GFP and RFP expression cassettes on their respective backbones. Media was changed within 24 hours of transfection. The following day, cells were sorted on a BD FACSAria for double-positive (GFP+/RFP+) expression to enrich for cells that had taken up both plasmids. Sorted cells were plated into a 6-well plate and expanded for one week. For clonal isolation, 2,000 cells were plated per 10 cm dish, grown for four days, and individual colonies were picked by hand into 96-well plates.

### G6-Well Cell Culture

Clonal lines in 96-well plates were maintained in N2B27 medium (as above). Cells were passaged every two days at a 1:6 split ratio using Accutase diluted 1:5 in PBS. Cell density was estimated by visual inspection under a microscope.

### Genotyping: Insert Copy Number Quantification

piggyBac insert copy number was estimated by TaqMan qPCR using absolute quantification with a standard curve. A TaqMan assay targeting the piggyBac inverted terminal repeats (ITRs) was compared to a single-copy genomic reference locus (Zeb2). Primers and probes were as follows: PB_TaqMan_Fwd, 5’-CACGTAAGTAGAACACGAAATAAC-3’; PB_TaqMan_Rev, 5’-GTGCTTGTCAATGCGGTAAG-3’; PB_TaqMan_Probe, 5’-/FAM/TGACTCACG/ZEN/CGGTTGTTATAGTTCA/3IABkFQ/-3’; Zeb2_Fwd, 5’-GGATGGGGAATGCAGCTCTT-3’; Zeb2_Rev, 5’-AGTGCGGCAGAATACAGCA - 3’; Zeb2_Probe, 5’-/HEX/TGATGGGTTGTGAAGGCAGCTGCACCT/ZEN/3IABkFQ/-3’.

### Insert Mapping by Ligation-Mediated PCR

Genomic insertion sites were mapped using a splinkerette-style ligation-mediated PCR approach adapted to the NEBNext system. Genomic DNA was fragmented using the NEBNext Ultra II FS DNA Library Prep Kit. NEBNext hairpin adapters were ligated to the fragmented DNA and subsequently cut with USER enzyme, generating non-complementary Y-shaped ends that suppress non-specific amplification from adapter-only fragments (analogous to hairpin suppression in traditional splinkerette PCR). A first round of PCR (20 cycles, 42 µl total volume) was performed using a primer to the 5’ piggyBac ITR (PB5R2, 5’-TCCAAGCGGCGACTGAGATG-3’) and a primer to the NEBNext adapter (NEBNEXTP7, 5’-GTGACTGGAGTTCAGACGTGT-3’). 1.5 µl of the first reaction was used as template for a nested PCR (20 cycles) with a second ITR-specific primer (PB_seq_read, 5’-TTTCCCTACACGCTCTTCCGATCT-3’)The nested PCR product was cleaned with AMPure XP beads at a 0.63x ratio. Indexing PCR was performed for 6 cycles using i7 and i5 primers, followed by a 0.7x AMPure XP bead cleanup. Libraries were pooled and sequenced. Only 5’ ITR junctions were mapped.

### RNA Extraction and Sequencing

RNA was extracted from approximately 100,000–200,000 cells per sample (one well of a 48-well plate) using TRIzol reagent, followed by purification with the Zymo Direct-zol RNA 96-well kit. RNA was eluted in nuclease-free water, quantified by Qubit fluorometer, and stored at −80°C. A total of 724 samples were submitted to Novogene across three sequencing batches. Novogene performed library preparation, quality control including RIN assessment (all samples passed), and sequencing. Samples were sequenced on an Illumina platform at 150 bp paired-end, yielding approximately 6 GB of raw data per sample (∼20 million read pairs).

### Luciferase Reporter Assay

F121 ESCs were plated at 14,000 cells per well in a 96-well plate one day prior to transfection. Cells were transfected with 200 ng of total DNA per well using Lipofectamine 2000 at a 3:1 ratio of Lipofectamine (µl) to DNA (µg). The transfection mix consisted of a firefly luciferase reporter plasmid (driven by the SV40 promoter with the candidate enhancer sequence) and a CMV-Renilla luciferase control plasmid at a 10:1 ratio (firefly:Renilla). Luminescence was measured 48 hours after transfection using the Promega Dual-Luciferase Reporter Assay System. Firefly luciferase values were normalized to Renilla luciferase to control for transfection efficiency.

### Computational methods

#### Insertion-site mapping

##### Sequence data and read structure

Splinkerette libraries were sequenced as paired 150-nt reads (Novogene). Read 1 begins in the piggyBac 5′ ITR and reads outward across the ITR– genome junction; read 2 begins at the splinkerette adapter and marks the opposite end of the sheared fragment. The i7 and i5 indices were retained in the read header throughout.

##### ITR extraction (stage 0)

Read pairs were selected with a multithreaded C extractor. Read 1 was matched against the 73-nt constant sequence immediately internal to the ITR terminus (CGCGCTATTTAGAAAGAGAGAGCAATATTTCAAGAATGCATGCGTCAATTTTACGCAGACTATCTTTCTAGGG) under a Levenshtein budget of 20 edits over 73 nt; mispriming products require ≥52 edits, so the threshold is permissive without admitting off-target priming. Matching pairs were written out with the 73 nt removed from read 1. The 73-nt trim point ends immediately 5′ of the TTAA target-site duplication, so the TTAA remains on the genomic side of the trimmed read and can be used as an internal check on junction placement rather than being consumed by trimming.

##### Trimming and quality filtering (stage 1)

Extracted pairs were processed in 16 interleaved shards by a streaming Python script (pb_prep.py), with shard outputs concatenated in order so that R1 and R2 remain in register; read-name synchrony was asserted on every record. Reads containing donor-plasmid backbone motifs were discarded. Each read was truncated at the first of: an exact 8-mer match to any TruSeq adapter arm or the read-2 adapter; the reverse complement of the terminal 16 nt of the ITR (read 2 only, marking read-through into the ITR); or a run of ≥8 G, which on two-colour chemistry represents absence of signal rather than sequence. Read 1 was required to retain ≥30 nt at mean Phred ≥25 after trimming; pairs failing this were dropped. Read 2 failing the same criteria was replaced by a length-30 all-N placeholder rather than discarded, so that the pair is retained for junction calling while being explicitly marked as carrying no usable fragment-end information. The full read header, including the i7+i5 index pair, was preserved.

##### Alignment and junction calling (stage 2)

Trimmed pairs were aligned to mm10 (UCSC) with bowtie2 v2.5.5 in local mode (--local -X 2000 --dovetail --sam-no-qname-trunc, 64 threads). The SAM stream was piped directly into the junction caller (pb_call.py) and never written to disk; the reference was held in memory so that genomic context could be retrieved without per-read indexing calls.

For each read pair, the junction anchor is the genomic coordinate of the 5′-most base of read 1, corrected for soft-clipping and for strand. A junction was called if read 1 aligned with MAPQ ≥30. Pairs in which only read 2 was confidently placed were retained as approximate sites, with the position estimated by subtracting the strand-specific median R1–R2 anchor offset, calibrated per run from the first 200,000 concordantly aligned pairs; these are validated to a median error of 47 bp (99^th^ percentile error 280 bp, 100% within 1 kb) and are flagged and excluded from all position-dependent analysis. Reads were tiered as both mates mapped, R1-only with unusable R2, R1-only with unmapped R2, or R2-only approximate.

Twelve bases of reference context around each anchor were extracted and searched for TTAA at offsets 0 and ±1 to ±4. Junctions were classified as canonical (TTAA at offset 0), shifted (TTAA within ±4), non-canonical (no TTAA), anchor-inferred (anchor derived from a soft-clipped alignment), or not called (approximate sites). TTAA status is recorded but is never used as a filter, so that non-canonical integration and local misplacement can be quantified rather than silently removed. Output is a per-read junction table (coordinate, strand, tier, TTAA class, 12-bp context, index pair) plus a 50-bp-binned site summary.

#### Well-resolved analysis

##### Exact site keys

Sites were keyed as chromosome:position:strand:population, with the position shifted to the TTAA where one was resolvable and left at the raw anchor otherwise; populations are canonical, non-canonical and anchor-inferred. The 50-bp binning of the pipeline’s own site table was not used for any counting.

##### Deduplication

The first 30 bp of the trimmed read 2 was used as a fragment-end fingerprint, joined to each junction record by read name through a 500,000-record sliding window (alignment reorders only locally); unmatched records were counted rather than assumed absent. Within each site, fingerprints were collapsed at Hamming ≤2 by greedy star clustering: representatives are taken in descending read count and absorb only sequences within 2 of the representative itself, bounding cluster diameter. Single-linkage or union-find clustering was explicitly avoided, as the transitive closure chains genuinely distinct fragment ends in low-complexity regions. A pigeonhole block index (two 30-mers within Hamming 2 must share one of three exact 10-bp blocks) makes this tractable at scale. Deduplication precedes all counting.

##### Index-hop correction

Two independent corrections were applied, because they fail in opposite regimes. First, each distinct molecule observed in more than one well was assigned a parent well (the well carrying the most reads of that molecule); copies in other wells were counted as hopped in. As a check that the shared observations are hopping rather than fragment-end collision, the fraction of hopped copies differing from the parent by exactly one of the two indices was reported. Because most molecules are singletons, this catches a hopped copy only when the parent copy was also sampled; orphan hops therefore survive. Second, a ratio filter retained only wells whose primary-molecule count exceeded 1% of the site’s top well, which cuts through the ∼150-fold abundance cliff between a real well and its hop tail.

##### Construct-borne loci

Sequence carried on the construct is present in every cell and therefore shows a flat well profile. Sites with ≥20 molecules across ≥8 wells and top-well dominance <0.40 were marked flat, flat sites within 3 kb were linked, and clusters spanning ≥20 sites were reported as candidate construct loci. The extended-cluster requirement distinguishes these from isolated flat positions, which are a separate artifact. Candidate loci were confirmed against the construct map before being blacklisted (±500 bp) from final counts.

##### Site classification and per-well typing

Surviving sites were classified using the two index axes separately, since well count and dominance alone cannot distinguish a construct locus from a clone associated with one pool. Where construct elements were supplied, per-well identity was called by a binomial test of each well’s element composition against the global element mix (the expectation for a well whose construct molecules were entirely hopped in), Bonferroni-corrected across wells. Purity was additionally tested for confounding with each index axis; where wells nest within perturbations, high per-well purity is an artifact of the design and is not evidence of perturbation-specific selection, which requires replication across independent pools.

##### Per-insertion provenance table

A final table was emitted with one row per well × site, carrying read and molecule counts, placeholder (no-usable-R2) counts, all-index read totals and per-axis polyG rates, molecule parentage (primary, hopped-in, ambiguous), and the ratio-filter outcome. Each site was assigned a confidence tier by testing n(s,c) ∼ Binomial(N(s), p(c)), where p(c) is the well’s share of the library — the appropriate null because both index hopping and mismapping distribute into a well in proportion to its depth. The test was run on deduplicated molecules and, failing that, on reads (yielding “confident” and “likely” respectively), with Bonferroni correction over the site–well pair count. Binomial upper tails were evaluated exactly as the regularized incomplete beta function via a modified Lentz continued fraction, so cost is independent of depth. A read-count-only fallback criterion for high confidence was deliberately not applied, since molecules are known at every depth here; applying it promotes deep sites lacking molecular support.

### RNA-seq analysis

#### Gene count processing

Gene-level read counts were obtained from featureCounts (v1.5.3) output files, with reads partitioned into three categories: genome 1, genome 2, and unassigned (*65*). Reads were aligned with STAR (v2.7.10b) to a dual N-masked mm10 reference generated by SNPsplit (v0.6.0) in two-genome mode, in which all known SNP positions distinguishing 129S1/SvImJ and CAST/EiJ from the C57BL/6J reference (Sanger Mouse Genomes Project, v5, dbSNP142) were replaced with N. Aligned reads were then partitioned by SNPsplit using strain-specific SNPs: reads overlapping one or more informative SNPs were assigned to the 129 or CAST genome, while reads overlapping no informative SNP or carrying conflicting allelic information were classified as unassigned. Each gene’s annotation lists coordinates for all of its exons. From these, the transcription start site (TSS) was defined as the start of the first exon for genes on the plus strand and as the end of the last exon for genes on the minus strand. For each category, counts were assembled into a samples × genes matrix. The genome 1 and genome 2 matrices were then reordered so their samples matched the order of the unassigned matrix. For descriptive purposes, counts were also converted to within-sample proportions by dividing each sample’s counts by its total count. All statistical modelling, however, used raw integer counts.

#### Negative binomial model of allele-resolved expression

Raw read counts were modelled with a gene-wise negative binomial (NB) generalized linear model. Counts were not normalized beforehand, which keeps their integer structure. Sequencing depth entered the model as an offset instead. Reads were split by allele, so each sample *c* is one cell line × allele row: allele 1, allele 2 or unassigned (U). For gene *g* in sample *c*, the count *Y_gc_* was modelled as

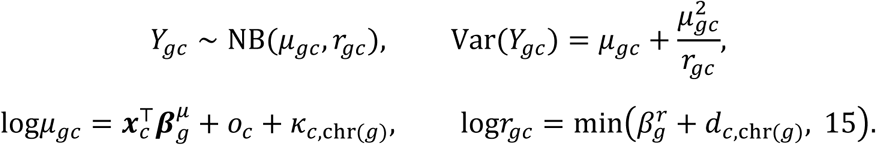

The covariate vector ***x****_c_* contains:

- an intercept;
- indicators for allele 2 and U (reference: allele 1);
- batch group, with three levels (batch1,batch2,batch3);
- perturbation type, with levels N, I, D and K (reference: S) referring to the inserted enhancer (Nanog:N, Ifitm:I,Sox2 ΔCTCF:D, Klf4:K);
- five latent *trans* factors (see *Latent trans factors*).

All genes on a chromosome share the same design matrix. Columns with no non-zero entries on a chromosome were dropped. The NB size parameter (inverse dispersion) has only a gene-specific intercept *β*_g_*^r^*. It is shifted by a fixed offset *dc*,chr for each sample and chromosome, and log*r* is capped at 15, which is effectively the Poisson limit. The library-size offset is *o_c_* = log*N_c_*, where *N_c_* is the sample’s total count across all genes, floored at 1. *κ_c_*_,chr_ is a copy-number offset. Both offsets have coefficients fixed at 1, so exp(***x****c*^T^***β****^μ^g*) is the expression rate per unit of sequencing depth at baseline copy number. With *pgc* = *rgc*/(*r_gc_* + *μ_gc_*), the log-likelihood of one observation is

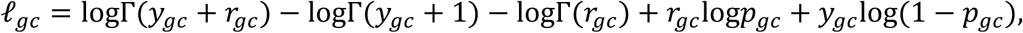

where log*p_gc_* and log(1 − *p_gc_*) are computed in log space for numerical stability.

#### Training mask and held-out observations

Model fitting used only observations in a training mask *M_gc_*. Observations that might carry an insert effect therefore could not influence the background model. Two kinds of region were masked (*M_gc_* = 0). First, genes whose transcription start site (TSS) lay within 2 Mb of any candidate insert in a line were masked in all three allele rows of that line. This step used a recall-oriented insert set. Second, each scored insert was matched to two control regions. A control region is the same locus in a randomly chosen other line that has no insert within 2 Mb. Genes within 500 kb of the locus were masked in the control line. Insert and control regions are therefore both scored out of sample.

The following were excluded from the analysis:

- lines without an insert map;
- one line with a known sequencing error;
- genes with fewer than 10 training reads, which were not fitted;
- for each gene, any allele row block with fewer than 5 training reads, which has no finite maximum-likelihood estimate (for example, genes without allele-informative SNPs).

Predicted genes (Gm-prefixed) and pseudogenes (-ps suffix) were not scored. It was confirmed programmatically that every scored observation was held out of training.

#### Latent trans factors

Five per-line latent factors were included as mean covariates. Each factor takes the same value on all three allele rows of a line. To estimate the factors, the model without factors was first fitted to all lines. For each line and each expressed gene (median fitted training reads per line ≥ 10), the residual log((*K* + 1)/(*M* + 1)) was computed. Here *K* and *M* are the observed and fitted training reads, summed over the line’s allele rows. Residuals were standardized per gene. The factors were the leading components of an iterative singular value decomposition (soft-impute), which filled held-out entries from the current low-rank reconstruction. Factor scores were scaled to unit variance.

### Copy-number and dispersion offsets

Coefficients were fitted chromosome by chromosome in three passes.

#### Baseline fit

The model was fitted with the library-size offset only.

#### Copy-number offset

For each sample and chromosome, the copy-number offset was the shrunk log ratio of observed to expected training reads:

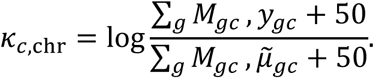

Here μ*_gc_* is the baseline-fit mean, capped at 10 × (the gene’s maximum training count + 1). Predicted genes and pseudogenes were left out of both sums. The model was then refitted with *κ* as a fixed offset.

#### Per-sample dispersion shift

For each sample and chromosome, *d_c_*_,chr_ was chosen to maximize the NB log-likelihood of the training observations under the current fit, plus a standard Gaussian log-prior −*d*^2^/2. The maximum was found by grid search over [−3,3] in steps of 0.05 and refined by parabolic interpolation. The model was then refitted with *d* as a fixed offset on log*r*.

### Parameter estimation

The objective was the negative log-likelihood over training observations plus an L2 penalty on non-intercept coefficients: 10^−2^∑(*β^μ^*)^2^for the mean and 10^−4^∑(*β^r^*)^2^for the size. The penalty keeps coefficients finite when an indicator column perfectly separates the data. Non-intercept columns were mean-centred during fitting, so the intercept represents the gene’s average expression level. Coefficients were transformed back to the original design afterwards. Coefficients were initialized by the method of moments, with all non-intercept coefficients set to zero. The mean intercept was set to 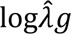, where 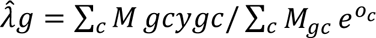 is the gene’s pooled rate. The excess dispersion

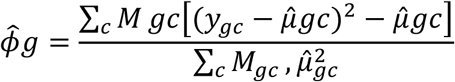

was clipped to [*e*^−14^, 100], and the size intercept was set to −logϕ*_g_*. Optimization used Adam in PyTorch (v2.13.0) in double precision, with (*β*_1_, *β*_2_) = (0.9,0.999) and a separate learning rate for each gene, starting at 0.05. It ran in stages:

1. the mean coefficients, with the size fixed;
2. the size, with the mean fixed;
3. steps 1 and 2 again;
4. all coefficients jointly.

Within each stage, progress was checked every 50 iterations. If the best loss in a window improved on the previous best by less than a relative 10^−9^, that gene’s learning rate was halved. A gene was considered converged once its learning rate fell to 10^−4^ or below and no coefficient moved by more than 10^−4^over a window. Each stage was capped at 20,000 iterations. Genes are independent and were fitted in chunks of 4,000. All fitted genes converged.

### Whole-line cross-fitting

Lines were split into three folds, stratified by batch group × perturbation type. For each fold, the gene-level coefficients were fitted without any of that fold’s lines. Every line was then scored with the fit that excluded it. The per-sample offsets *κ* and *d* can only be estimated from a line’s own data. For held-out lines, these offsets were therefore estimated from the line’s training observations only, never from its scored windows.

### Significance of held-out observations

Each held-out observation was scored by its upper-tail probability under the fitted NB distribution:

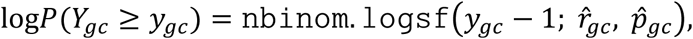

computed with SciPy (v1.18.0). The fitted parameters *μgc* and *rgc* include all fixed offsets (library size, copy number and dispersion shift). No observation-specific term was fitted. We used the inclusive tail *P*(*Y* ≥ *y*), hence the *y* − 1 argument, because the exclusive tail *P*(*Y* > *y*) is anti-conservative at low counts. Each region (an insert or control locus in one line) was scored separately at radii of 50, 100, 250 and 500 kb. The region statistic was the smallest p-value among all tested gene × allele observations with a TSS within the radius, Šidák-corrected for the number of observations *n*: 1 − (1 − *p*_min_)*^n^*. Significance thresholds were calibrated on the hit rate in matched control regions (for example, the threshold that gives a 1% control hit rate), rather than by a per-observation false discovery rate procedure.

### Effect size (Δ expression / shrunk log fold change)

The expression effect at each held-out observation was estimated as a ridge-penalized NB log fold change, computed after the model fit. It was not used in the significance test. For each insert × gene pair, the effect was computed on the same allele row as its p-value. This was the candidate row (the insert’s haplotype allele or U) with the smallest p-value. All coefficients and offsets were held at their cross-fitted values. The fitted mean *μ_gc_* and size *r_gc_* were therefore fixed, and they include the library-size and copy-number offsets and the dispersion shift. A single multiplicative shift on the mean was then estimated for the observation:

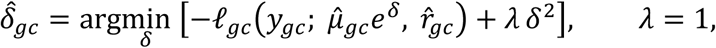

where l*_gc_* is the NB log-likelihood defined above. As in the model fit, 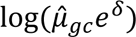 and 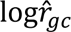 were clamped to [−20,20]. The penalty shrinks 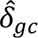 toward zero. Shrinkage is strongest at low expected counts, where the raw ratio 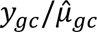 is noisy. The problem for each observation is one-dimensional and strictly convex. It was solved by Newton’s method starting from *δ* = 0. Steps were capped at magnitude 1 so that iterates could not overshoot past the clamp, and iteration stopped at a step-size tolerance of 10^−10^. 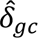 is on the natural-log scale, and positive values indicate expression above the background prediction. Δ expression in the figures is 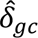.

### Specific analyses

#### Insert and cell line filters

Lines were excluded if the allelic copy-number ratio on the insert’s chromosome deviated from 1 by more than 0.25, and a fixed list of QC-failed lines was removed (“karyotype-clean”). An insert was isolated if no other insert of the same line lay within the stated distance; an insert alone on its chromosome counted as isolated. Insert confidence means either a DNA-based confidence score or RNA confirmation by reads spanning the construct–genome junction. All distances are signed relative to insert orientation. The table gives the settings per analysis; figure descriptions below state only departures from it.

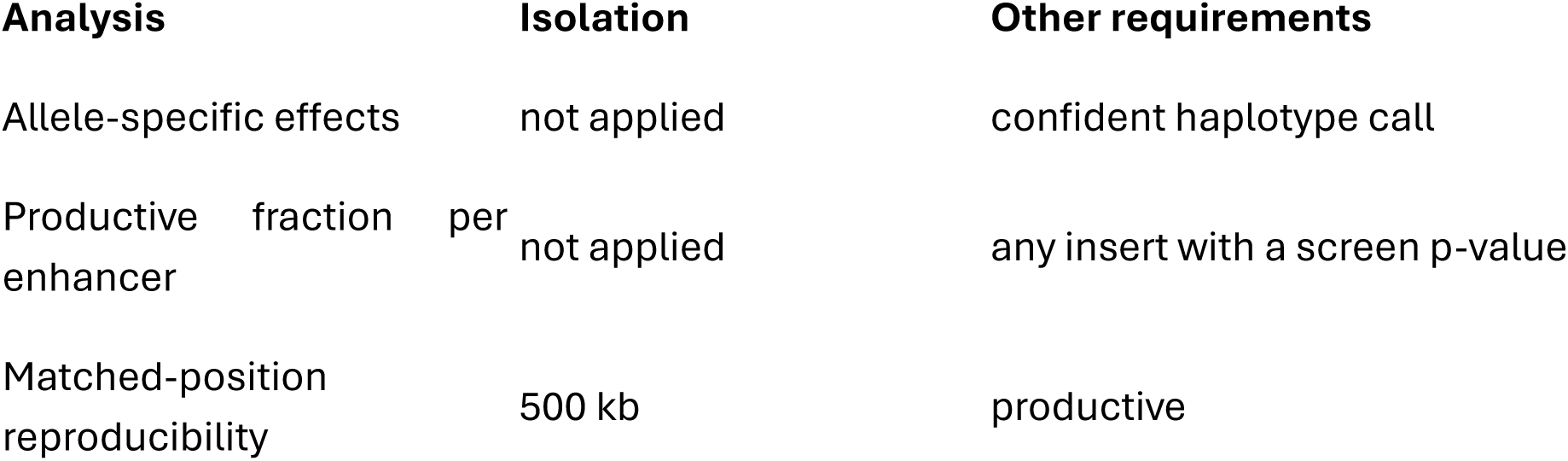
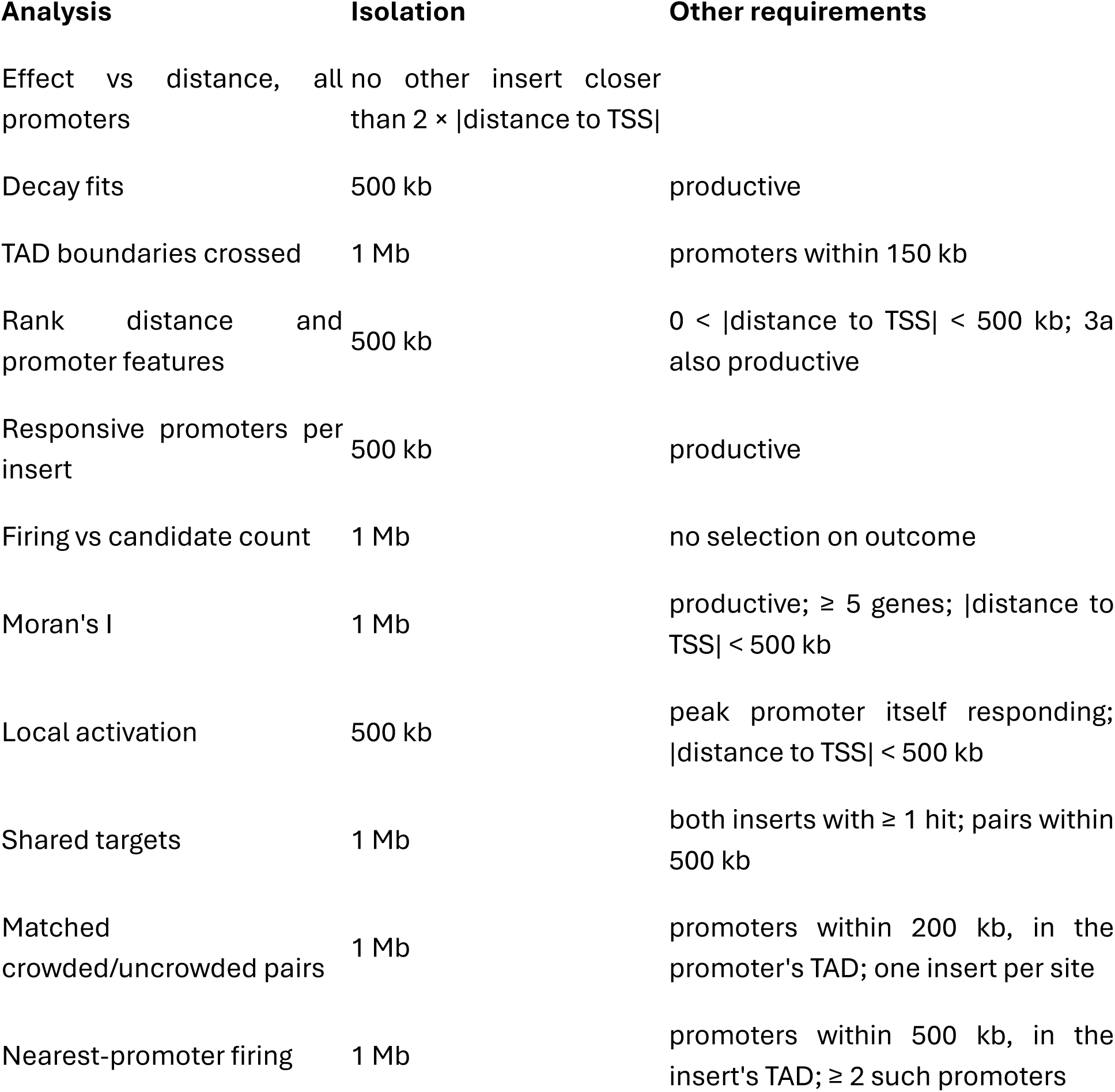

#### External datasets

Micro-C was balanced and used at 1 kb resolution unless otherwise stated. A promoter is in the insert’s TAD when no boundary lies between the insert and its TSS. Donor enhancer signal [1b] was Gaussian-smoothed (σ = 20 bp), with each track’s y-axis shared across the four regions.

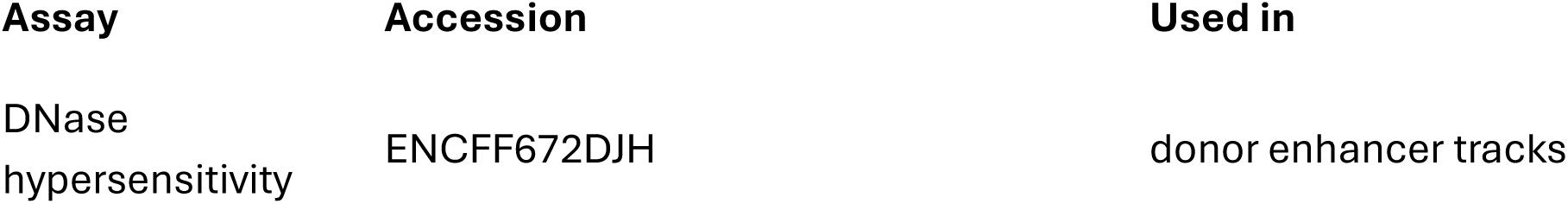
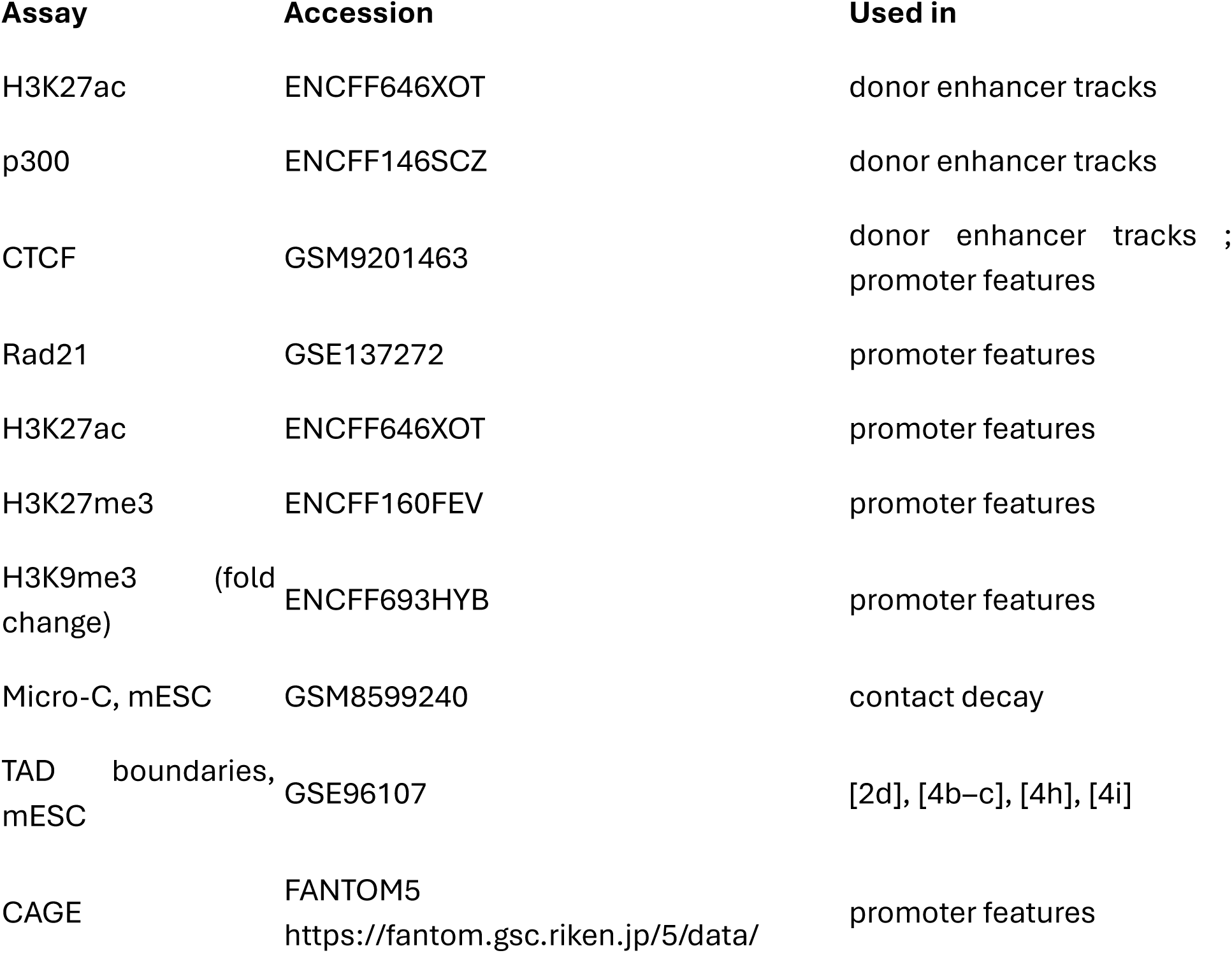

#### Allele-specific analysis

Sites needed a confident haplotype call: ≥ 10 insert-mapping reads and ≤ 5% of reads assigned to the other haplotype by SNPsplit (17,006 of 29,954 sites). Delta expression was fitted separately for each allele. For genes within 200 kb whose insertion allele responded, insertion- and opposite-allele delta expression were plotted as a Gaussian KDE against both alleles of all promoters within 200 kb: 7,576 of 70,724 proximal site × gene pairs at 5,542 sites, against a distal background of 1,055,888 allele tests. Insertion versus opposite allele was tested by how often the opposite allele was also significant by a one-sided binomial test against the distal rate (|distance to TSS| ≥ 200 kb, either allele).

#### Reproducibility across independently derived inserts

Inserts in different lines lying within 100 bp of each other were compared, carrying either the same [1f] or different [1g] enhancers. For each gene within 500 kb of both inserts, Delta expression was compared by Pearson’s r. Same enhancer: 1,966 pairs, 2,386 inserts, 28,844 gene comparisons. Different enhancer: 142 pairs, 203 inserts, 2,002 comparisons.

#### Distance decay

The relationship between insert–promoter distance and Δ expression was characterized by fitting two models to bin-averaged Δ expression across all promoters (n = 141,850; 30 logarithmically spaced distance bins). First, a four-parameter sigmoidal curve was fitted by nonlinear least squares (scipy.optimize.curve_fit), yielding R² = 0.98. Second, a ceiling-only model was fitted in which Δ expression decays as a power law of distance (exponent *k* = 1.18), capped at each promoter’s dynamic range, defined as the difference between the 95th and 5th percentiles of that gene’s FANTOM5 CAGE expression across samples (R² = 0.98). For comparison, a Hill curve was fitted to Micro-C contact frequency between flanking regions at matched distances.

#### TAD boundaries

Boundaries crossed between insert and promoter were counted for all promoters within 150 kb and for regulated promoters at ln p < −10. *n* = 100,581 promoters (all) and 4,631 (regulated).

#### Promoter rank

Rank is the gene’s order outward from the insert on its own side, nearest ranked 1, among genes with a screen measurement. Responding promoters at productive inserts were counted by rank, normalized per enhancer and pooled, with ranks above 10 combined. *n* = 9,042 responding promoters at 6,724 inserts (Sox2 1,224; Nanog 2,407; Ifitm 535; Klf4 4,568; Sox2 ΔCTCF 308). The example locus is Trit1/Mycl at a Nanog insert, ±150 kb, with arcs at ln p < −10.

#### Promoter features

The 30 nearest genes on each side of each insert were classified as significant (log p < −10); skipped (not significant, but nearer the insert than a significant promoter on the same side); or non-significant (all other promoters of inserts with at least one significant promoter). Features are means over TSS −1 kb/+0.5 kb, strand-oriented, stratified by insert distance (0–10, 10–100, 100–500 kb): CTCF, Rad21, H3K27ac, H3K27me3 and H3K9me3 fold-change ChIP signal; promoter GC content; FANTOM5 CAGE Gini coefficient and 95th–5th percentile range for the gene’s dominant peak; and basal expression. Missing values were excluded, CAGE measures being absent for genes without a peak. Violins include box-and-whisker summaries.

#### Responsive promoters per insert

Promoters at ln p < −10 within 500 kb were counted per insert, counts above 10 combined. *n* = 6,724 inserts.

#### Firing against candidate count

Mean promoters firing against candidate promoters, in the insert’s TAD [4b] and within 150 kb of it [4c]. Inserts were not selected on outcome, so inserts that activated nothing count as zero.

#### Spatial clustering

Activation was ln p divided by the region’s minimum ln p, on a 0–1 scale. Moran’s I used weights *w_ij_* = exp(−|*TSS_i_* − *TSS_j_*|/bandwidth); mean I is shown with its 30th–70th percentile band at window sizes from 5 to 500 kb. *n* = 5,501 regions. Here TSS means distance to TSS.

#### Local activation

Median delta expression against distance from the region’s most activated promoter, compared with a random non-peak promoter matched on distance to the insert. Bands are 95% bootstrap CIs on the median. n = 1,879 regions, of 3,161 with ≥ 10 promoters.

#### Shared targets

For insert pairs within 500 kb, the shared fraction was 2·|shared hits| / (hits₁ + hits₂), grouped by inter-insert distance and by the number of promoters between them (0, 1, 2, ≥ 3). Means with 95% bootstrap CIs; cells with fewer than 10 pairs were dropped. *n* = 13,683 pairs.

#### Matched crowded and uncrowded sites

The same promoter was compared at two insert sites in its own TAD: nearest promoter at one, with at least two promoters closer at the other. The crowded site had to be the closer of the two by no more than 1.5×, so the distance difference works against the result. A4gnt is shown as an example. P(fires) was averaged per promoter, one vote per promoter, with 95% bootstrap CIs over promoters and a two-sided sign test on discordant promoters. *n* = 110 promoters, 148 site pairs.

### RbP modeling

#### Row selection

Training rows were selected by the filters below.

| Filter | Setting |
| --- | --- |
| Gene window | Distance to the insert was less than 1 Mb |
| Site isolation | No other surviving insertion within 1 Mb. |
| Site quality | High confidence insertion |
| Group responded | There was at least 1 strong responding promoter ($\ln p < -20$) within 500kb of the insert |
| Karyotype QC | Not on a karyotypic abnormal chromosome |
| Genes | GENCODE-ordered protein-coding and lncRNA genes; -ps and Gm\d pseudogenes dropped |

#### Labels

Each gene–insertion pair was assigned an ordinal severity label by binning its log survival-function value (equivalent to the ln p-value of the per-gene negative binomial test) at thresholds of −5, −10, and −20, yielding four ordered classes: 0 (ln p > −5), 1 (−10 < ln p ≤ −5), 2 (−20 < ln p ≤ −10), and 3 (ln p ≤ −20). A pair was called a hit at the most stringent threshold, ln p ≤ −20.

#### Promoter features

Six per-gene promoter features form the promoter feature matrix (49,997 genes).

| Feature | Definition |
| --- | --- |
| gc_core | GC content of TSS ± 50 bp |
| dinuc_TA | TA dinucleotide frequency over the strand-aware promoter window [TSS – 1000, TSS + 500) |
| p95_p5_range | Cross-sample 95th–5th percentile range of $\log(\text{TPM} + 1)$ over the FANTOM5 CAGE panel, at the gene's highest-expressed peak |
| gini | Gini coefficient of the same CAGE distribution |
| H3K27ac | Mean mESC ChIP bigwig signal over [TSS – 1000, TSS + 500) |
| H3K27me3 | Mean mESC ChIP bigwig signal over the same window |

All six are promoter measurements of the gene alone. The signal matrix is indexed by gene_id, so any column placed there is constant across insertions by construction, and any quantity that varies with the insertion must travel on the pairs frame instead.

#### Contact

Contact is ins_contact_DMSO, the predicted contact of the insertion allele taken from the contact prediction model, in log space.

### Model

#### Raw regulatory score

For enhancer–gene pair i in group g, the regulatory score is

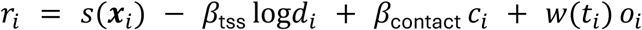

where the terms are defined as follows. Promoter term *s*(***x***). A three-layer MLP (32 hidden units; Linear → LayerNorm → GELU → Dropout) maps the six standardized promoter features to a scalar. The dropout probability of 0.707 was selected via an Optuna hyperparameter sweep. This high dropout likely prevents the model from memorizing the training data improving generalizability. Distance term *β*_tss_log*d_i_*. A learned coefficient on the log-distance from the enhancer to the TSS, penalizing distal pairs. Contact term *β*_contact_ *c_i_*. A learned coefficient on the standardized predicted 3D contact frequency. Orientation term *w*(*t_i_*) *o_i_*. A perturbation-type-specific weight *w*(*t*) — one learned scalar per type, zero-initialized — multiplied by a signed oriented log-distance:

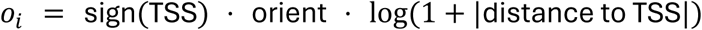

#### Competition

The raw regulatory score is divided by a learned temperature T = softplus(θ) + 0.1, clamped to ±10, and normalized within the group:

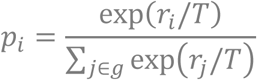

#### Severity heads

Three independent binary heads (2 layers, 64 hidden units, dropout 0.367) predict the cumulative thresholds −ln p ≥ 5, 10 and 20 from the promoter features as well as the partition of the enhancer output received.

#### Standardization

A single InputScaler, fitted on the training split alone, standardizes the promoter feature matrix, log1p|distance to TSS|, the contact column, and the oriented TSS.

### Network training

#### Objective

The only loss is the ordinal loss on the severity heads: binary cross-entropy on the three cumulative thresholds, with per-threshold class weights n_below / n_above capped at 50, averaged over thresholds and over rows.

#### Optimization

| Setting | Value |
| --- | --- |
| Split | Insertions on chr1 and chr2 held out, by insertion chromosome |
| Optimizer | AdamW, learning rate $4.1900 \times 10^{-3}$, weight decay $2.1826 \times 10^{-4}$ (0 on plateau parameters) |
| Schedule | Cosine annealing over 30 epochs |
| Batch | 32 competition groups, concatenated with group ids |
| Gradient clipping | Global norm 1.0 |
| Seeds | 42, 1, 2; numpy, torch and CUDA all seeded |
| Initial $\beta$ | tss 4.955 , contact 2.514, offset 1.731 |

Hyperparameters were selected by an Optuna sweep using chr3 and chr4 as the optimization target.

#### Ablation protocol

The term is gated to zero and its parameters frozen, so an ablated arm is architecturally identical to the full model.

The No competition arm replaces the grouped softmax with sigmoid(a_i_/T) for every candidate.

#### Transfer to the K562 CRISPRi benchmark

The mouse checkpoint was applied to human data without retraining.

#### Rows and labels

Pairs were taken from EPCrisprBenchmark_combined_data.training_K562.GRCh38.tsv.gz (https://github.com/EngreitzLab/CRISPR_comparison/tree/main; 10,356 pairs), with one group per tested element. Groups were kept when they contained at least one Significant pair with EffectSize < 0 — the analogue of RbP’s region_min condition, i.e. the element did something — and at least two candidate genes, leaving 1,069 pairs and 292 positives across 231 element groups.

#### Mapping onto the model

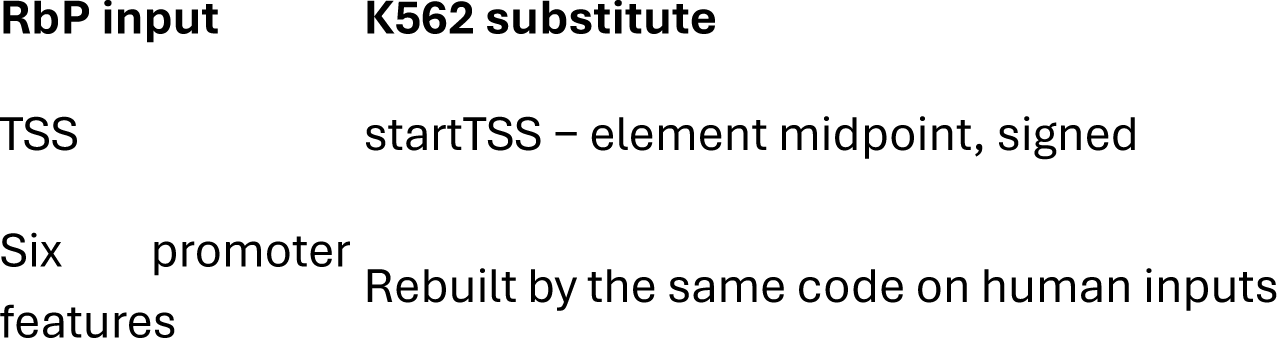
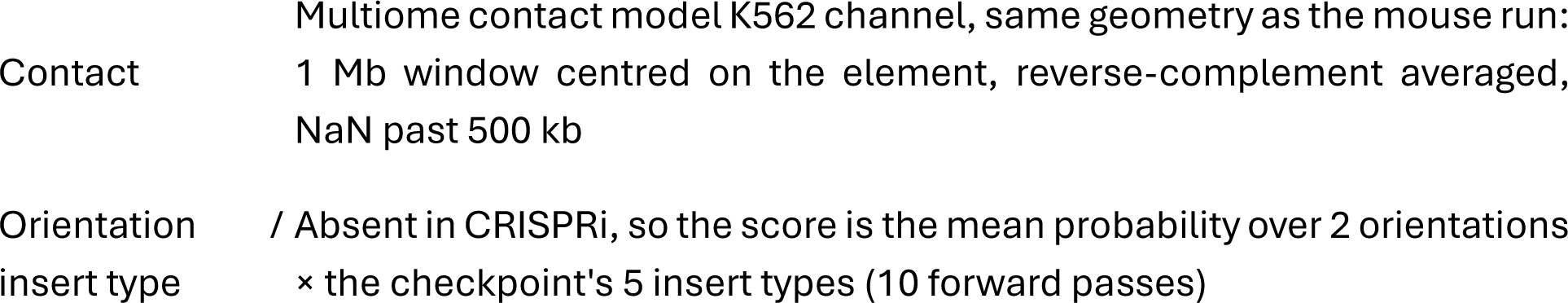

Human features use the identical feature code pointed at human inputs: hg38 sequence for gc_core and dinuc_TA over the strand-aware [TSS − 1000, TSS + 500) window, with strand from GENCODE v46 and the TSS from the benchmark itself; the 1,829-sample FANTOM5 human CAGE panel (hg19 peaks lifted to hg38, each peak assigned to the nearest GENCODE TSS within 200 bp, highest-mean peak per gene) for p95_p5_range and gini; and K562 ENCODE H3K27ac (ENCFF381NDD) and H3K27me3 (ENCFF242ENK) for the two marks, preserving the mouse pipeline’s window and its NaN-on-gap behaviour.

#### Quantile mapping

K562 signals are quantile-quantile mapped to the mouse value at its own rank percentile: the six signal features onto the 49,997-gene mouse signal matrix, and contact onto the training frame’s ins_contact_DMSO.

### Comparison methods

#### ENCODE-rE2G: published code and published models

All rE2G predictions were produced by running the published rE2G code on the published rE2G models, without modification. The application step is X = log(|features| + 0.01); model.predict_proba(X)[:, 1] and only models/<name>/model.pkl and models/<name>/feature_table.tsv are read.

Three published models are used, one for each application below. The published K562 predictions use rE2G-extended, a 45-feature logistic regression with no penalty. The mouse insertion screen uses the 8-feature atac_h3k27ac_intact_hic model. The mESC enhancer knockouts use dhs_megamap, which additionally requires promoter accessibility as an input feature.

#### Published K562 predictions

Predictions were taken from the ENCODE portal as element–gene-link beds. Their elements are the pipeline’s own candidate regions and do not share coordinates with the CRISPRi tested elements, so, following EngreitzLab/CRISPR_comparison, each CRISPRi pair takes the maximum score over predicted links that name the same target gene and overlap the tested element.

#### Zero-shot and retrained rE2G on the mouse insertion screen

rE2G’s eight features were built for every (insertion, gene) pair, with the insertion site playing the role of the element.

Element activity. All inserted enhancers are treated as equally active: A_insert is a single constant for every insertion, the median endogenous candidate activity, so the construct is a typical enhancer.

- Contact. rE2G’s contactFrequency is a positive, power-law-scaled contact that the model logs, whereas ins_contact_* is already in log space, so it is exponentiated back to linear first. A missing contact is filled with the mean linear contact of its distance bin, so its deviation from expectation is exactly 1 and contactFrequency falls back to the power law, in accordance to ABC’s documented fallback. Contact-observed rows are also reported separately.
- ABC denominator. 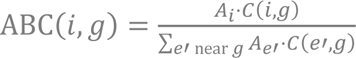, with the numerator from the predicted insertion-allele contact and the denominator over endogenous candidate elements using the power-law fallback C = d^γ.
- Retrained variant. This keeps rE2G’s architecture, feature set and transform and matches RbP on everything else: the same rows, the same chr1 + chr2 holdout, the label ln p < −20, the same contact column, and the cell-line exclusion list.
- Group reconditioning. Because RbP’s softmax normalises within a group, rE2G probabilities are also reported reconditioned as *p_i_*/(1 − ∏*_j_*(1 − *p_j_*)) over the insertion’s genes, treating pairs as independent.

#### AlphaGenome, K562 — published predictions for this benchmark

The K562 AlphaGenome column is DeepMind’s own published predictions for this benchmark (enhancer_gene_linking_e2g_predictions.feather, from the AlphaGenome evals bucket), joined onto the benchmark rows by (enhancer_id, gene_id): 10,353 of 10,356 pairs, column prediction, with output_type = RNA_SEQ and metric_calculator = EnhancerGeneLinkingAUCPR.

The method is the paper’s input × gradient scorer. A window is centered on the gene-body midpoint (GENCODE v46); the differentiated scalar is log(Σ RNA-seq signal across the gene’s exons), averaged over K562 RNA-seq tracks (EFO:0002067); and the element’s score is a Gaussian-kernel-weighted mean of |grad × input| centered on the element (2,400 bp window, σ = 300 bp), divided by the mean |grad × input| over the whole window.

#### AlphaGenome, mouse insertion screen

AlphaGenome was run on the insertion data the pipeline predicts 1 Mb windows for each insertion allele and for its matched reference allele, and derives 36 EFO track features plus 8 measured ENCODE columns. The reported score is a single derived feature: the log ratio of predicted H3K27ac at the target TSS between the insertion and reference alleles. Which provided the strongest single feature AUCPR for alpha-genome.

### Statistics and conventions

#### Metrics

AUCPR (sklearn.metrics.average_precision_score), pooled over pairs and stratified by |TSS| into < 10 kb, 10–100 kb and 100 kb–1 Mb (and > 1 Mb where that bin exists), with per-bin prevalence reported alongside, because positives are strongly distance-skewed and pooled AUCPR across a wide distance range flatters a distance heuristic.

#### Rank metrics

Within-element top-1 accuracy, mean reciprocal rank, hit percentage and nDCG are reported for the K562 element sets. Chance MRR is computed exactly rather than simulated, from 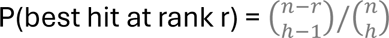 for a group of n candidates with h hits, and was verifiedagainst 4,000 permutations per group (0.5966 exact against 0.5972 permuted).

#### Reconditioning

To remove the advantage an unnormalized method gains from not being a softmax, each method’s score can be divided by its group total, *p_i_*/(1 − ∏*_j_*(1 − *p_j_*)) over the insertion’s genes, treating pairs as independent.

#### Uncertainty

Two kinds of error bar appear and are labelled as such.

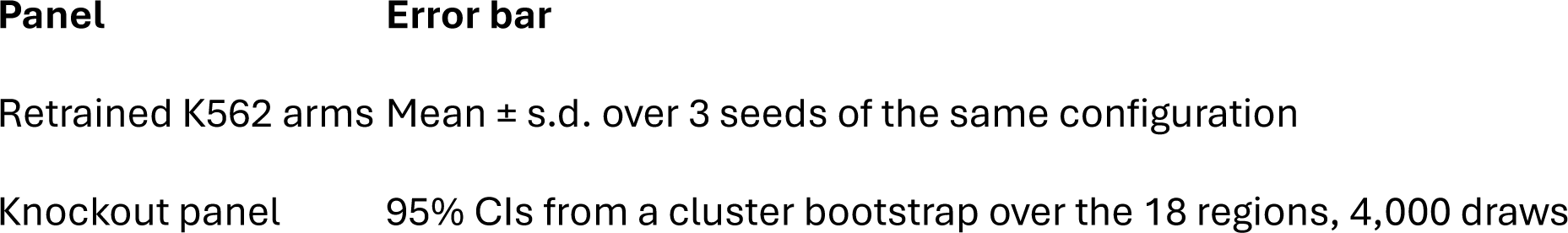

#### Gradient × input scoring of variant–gene pairs

Candidate regulatory variant–gene pairs were scored zero-shot using AlphaGenome, with the publicly released pretrained weights (google/alphagenome-all-folds) and the JAX reference implementation. Because the public AlphaGenome API provides only in silico mutagenesis for interpretation, input gradients were computed by backpropagating through the model’s forward pass using jax.grad. The model accepts a one-hot-encoded DNA sequence as input, which is differentiable.

#### Scalar objective

For each variant–gene pair we constructed a 1,048,576 bp (2²⁰) input window on GRCh38, centered on the midpoint of the gene body as defined by the GENCODE v46 basic annotation. The scalar objective differentiated was the log of the predicted RNA-seq signal summed over the gene’s exons:

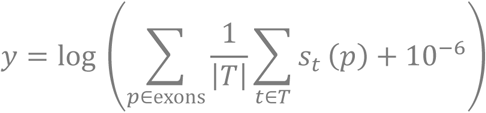

where *s_t_*(*p*) is the predicted 1-bp RNA-seq activation for track *t* at position *p*, *T* is the set of all AlphaGenome RNA-seq tracks for HFFc6 fibroblasts (ontology term EFO:0009318), and exonic positions are the union of the gene’s annotated exons (overlapping exons merged) clipped to the input window.

#### Variant score

The gradient of *y* was computed with respect to the one-hot reference sequence ***X*** ∈ {0,1}*^L^*^×4^ and formed a per-position attribution

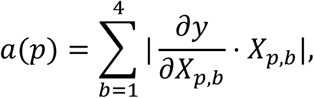

corresponding to the absolute gradient × input at the reference base. The score assigned to a variant at position *v* was the ratio of a Gaussian-weighted local mean to the global mean attribution:

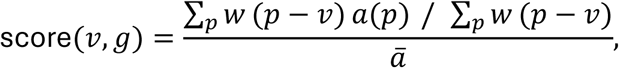

where *a* is the mean of *a*(*p*) over the full 1-Mb window, and *w* is a truncated Gaussian kernel with *σ* = 300 bp and support |*d*| < 1,200 bp:

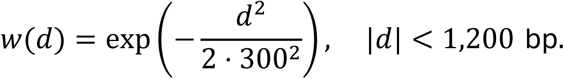

The score uses only the reference sequence, not the alternate allele, and measures how strongly the local sequence context of the variant contributes to the model’s predicted expression of the candidate gene. Because the window is centered on the gene rather than the variant, each variant–gene pair requires its own forward and backward pass. Pairs whose variant fell outside the input window (i.e., the gene-body midpoint was more than approximately 524 kb from the variant) received a score of zero and were retained in all evaluations; the same applied to pairs with no annotated exon in the window.

#### Validation

The implementation was validated on the K562 CRISPRi enhancer–gene benchmark (EngreitzLab CRISPR_comparison, training_K562; 10,356 pairs, 471 positives), using K562 RNA-seq tracks (EFO:0002067) and centering the Gaussian kernel on each tested element.

#### Evaluation on GTEx fibroblast eQTLs

Every candidate pair in the GTEx v8 cultured-fibroblast eQTL set was scored, comprising 9,448 variant–gene pairs, 1,663 variants, 4,187 candidate genes, and 1,673 targets. A pair was labelled positive if the variant is a significant eQTL for the gene (is_target). Performance was measured as the area under the precision–recall curve (AUCPR), both overall and stratified by absolute variant–TSS distance (0–10 kb, 10–100 kb, 100 kb–1 Mb). The 100 kb–1 Mb bin effectively covers 100–500 kb, because candidate genes were limited to ±500 kb of the variant. For comparison with the ENCODE-rE2G HFFc6 DNase-only model (ENCFF773XUW), the primary comparison was restricted to the 1,214 pairs (233 positives) whose variant lies within an rE2G candidate element linked to that gene. The 1,214 pair set is the one reported.

#### Group reconditioning

RbP outputs a softmax distribution over each variant’s candidate genes, whereas the other methods score each pair independently. To place all methods on a comparable footing, each method is also reported reconditioned on the assumption that the variant has at least one target gene:

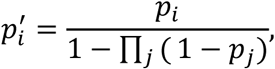

where *j* ranges over all candidate genes for that variant (the same group over which RbP’s softmax is computed) and pairs are treated as independent. RbP is not reconditioned, as its softmax is already conditioned on the group; its reconditioned score is therefore its raw score. ENCODE-rE2G scores were used as provided. Because gradient × input scores and inverse-distance scores are not probabilities, each was first mapped to *P*(target) using a logistic regression on log(score + 10^−3^). This calibration was fit out-of-fold with five-fold cross-validation grouped by chromosome; because the mapping is monotone, raw AUCPR is unaffected. Variants for which a method assigned a score of zero to every candidate gene retained a score of zero after reconditioning.

#### Beta globin locus predictions

The enhancer and Capture-C viewpoint were both centered on LCR HS3 at chr11:5,284,640 (hg38). The promoters within 500kb were considered.

#### Promoter Distance from HS3 (kb) WT group dProm group

|  |  |  |  |
| --- | --- | --- | --- |
| HBE1 | -14.7 | yes | yes |
| HBG2 | -29.9 | yes | yes |
| HBG1 | -34.8 | yes | yes |
| HBBP1 | -41.1 | yes | yes |
| HBD | -50.2 | yes | yes |
| HBB | -57.6 | yes | no (deleted) |
| OR51M1 | +99.2 | yes | yes |
| TRIM5 | +400.4 | yes | yes |
| TRIM22 | +405.1 | yes | yes |

#### Promoter features

Six features per promoter, in the order the scaler was fit. The table was computed once and shared by both conditions.

**Promoter features** Six features per promoter, in the order the scaler was fit. The table was computed once and shared by both conditions.
| Feature | Definition | Source |
| --- | --- | --- |
| gc_core | GC fraction, TSS $\pm 50$ bp | hg38 sequence |
| dinuc_TA | TA dinucleotide frequency, $-1,000/+500$ bp | hg38 sequence |
| p95_p5_range | 95th - 5th percentile of $\log(\text{TPM}+1)$ across samples | FANTOM5 CAGE |
| gini | Gini coefficient of $\log(\text{TPM}+1)$ across samples | FANTOM5 CAGE |
| H3K27ac | Mean signal -1,000/+500 bp around TSS, strand-oriented, quantile-mapped | HUDEP-2 ChIP-seq (GSM5267246) |
| H3K27me3 | Same window and mapping | GSM9452939 |

Because the model was trained on mESC marks, HUDEP-2 marks were quantile-mapped onto the training scale: promoter means for every gene in the hg38 table were converted to percentile ranks (clipped to [1e−6, 1−1e−6]) and each rank replaced by the matching mESC quantile for that mark. This preserves within-HUDEP-2 ordering.

Contact came from measured HS3-viewpoint Capture-C (WT: GSM5827251; dProm: GSM5827253), both normalized reporter bigwigs in hg38, converted to log observed/expected rows on the model’s contact scale. A 1 Mb window centered on the viewpoint, start-snapped to the 4 kb grid, was split into 250 bins of 4 kb. Each bigwig interval is one restriction fragment carrying a normalized reporter count; a fragment contributed value × (overlap / fragment length) to each bin it overlapped, so bins hold total reporter count rather than a length-weighted sum. Empty bins were set to 0. The expected-by-distance curve came from 4DN Hi-C (4DNFI9GMP2J8, 1 kb), built into a full matrix, block-averaged to 4 kb to match the contact model’s training pipeline, and read off at the viewpoint’s row; ε is the minimum of the rebinned expected matrix. Since Capture-C reporter units differ from balanced Hi-C units by an unknown factor, one scale factor α per profile was estimated as the median, across eight log-distance strata, of the per-stratum median log(obs/exp), using only bins >5 kb from the viewpoint. Contact per bin was

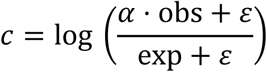

with bins within 5 kb of the viewpoint set to missing. Each promoter took the value of the 4 kb bin containing its TSS; all nine candidates are >14 kb from HS3, so all received finite values. Coarser bins would place HBG1, HBG2 and HBD together and give them identical contact.

#### Promoter deletion and readout

The deletion changed two inputs together: the dProm Capture-C profile replaced the WT profile, and HBB was removed from the competitive group. Everything else — gene set, feature matrix, promoter constant, model weights — was shared and computed once.

### Mouse ESC enhancer deletion benchmark

#### Candidate genes and labels

For each deletion, candidate target genes were all annotated genes with a TSS within ±500 kb of the deletion’s midpoint. For Etl4 the window was ±1 Mb. We then kept only genes expressed in mESC, meaning more than 1,000 summed reads across our RNA-seq samples

#### Scoring with RbP

We scored the deletions with the same RbP model used throughout the paper and did not retrain it on any knockout data.

##### Predicted contact

We ran the sequence-to-contact network (contact prediction model) on the 1 Mb reference mm10 sequence centered on each deletion midpoint. Bins were 4 kb (250 bins per window), and predictions on the forward and reverse-complement strands were averaged. Each candidate TSS was assigned the predicted contact between the enhancer’s bin and the TSS’s bin, read from the DMSO output channel. That channel is the contact input the RbP model was trained on (ins_contact_DMSO). For Etl4, TSSs outside the central window were reached with additional, shifted 1 Mb windows.

##### Gene-level features

Promoter features were the six signal features of the final model (GC content of the core promoter, TA dinucleotide frequency, p95−p5 dynamic range, Gini coefficient, H3K27me3 and H3K27ac) as the promoter term. All were taken from the same mESC signal matrix used for training.

##### RbP model

Each enhancer–gene pair was scored with the trained RbP. Four deleted elements (Sox2, Klf4, Nanog, Ifitm) are also inserted elements in the training screen, so they were given that element’s learned insert identity and + orientation. All other regions used the model’s default insert type. The ordinal severity head −20 bin was used for scoring.

#### Baselines

All baselines were scored on the same 268 enhancer–gene pairs as RbP.

- **1/distance.** The inverse of the absolute distance from the deletion midpoint to the TSS.
- **ABC.** Candidate elements were the top 150,000 mESC ATAC peaks (ENCODE ENCFF006ESP, ranked by peak height), each as 500 bp around its summit. Element activity was the geometric mean of ATAC and H3K27ac signal. For the deleted element itself, activity was measured over the full deleted interval. Contact was a power law in distance (γ = −1.024, floored at 1 kb). The ABC score was the element’s activity × contact, divided by the summed activity × contact of all candidates within 5 Mb of the TSS.
- **ENCODE-rE2G.** We applied the published DNase-only rE2G model (dhs_megamap) zero-shot. We computed its features on the mouse pairs, with predicted DMSO contact divided by its mean at that distance as the contact feature. The published model has no scaler, so the assay-dependent features (promoter accessibility, sumNearbyEnhancers, contactFrequency, ABC score) were quantile-normalized onto their distributions in ENCODE’s K562 predictions. Distances, counts and the ubiquitous-expression flag were left unmapped. Without this normalization rE2G scored at chance (AUCPR 0.101).

### XGBoost training

#### Target

The response variable was −*ln p*. It was discretised into six ordinal bins (0, 1, 2–5, 5– 8, 8–20, ≥20), and a gradient-boosted tree classifier (XGBoost) was fitted over all six bins with a multi-class softmax objective (multi:softprob). All quantities reported here use the predicted probability of the top bin (ln p ≤ −20), matching the label used in the RbP sweeps. The multi-bin ordinal regression acted as a multitask regularization step improving performance.

#### Training and early stopping

Chromosome 3 was held out as the validation set for early stopping. Training ran for up to 6,000 boosting rounds at a learning rate of 0.015 and stopped when validation AUCPR had not improved for 150 consecutive rounds. Trees were grown with the histogram method on a single GPU. Hyperparameters are given in Table 2.

**Table 2.**
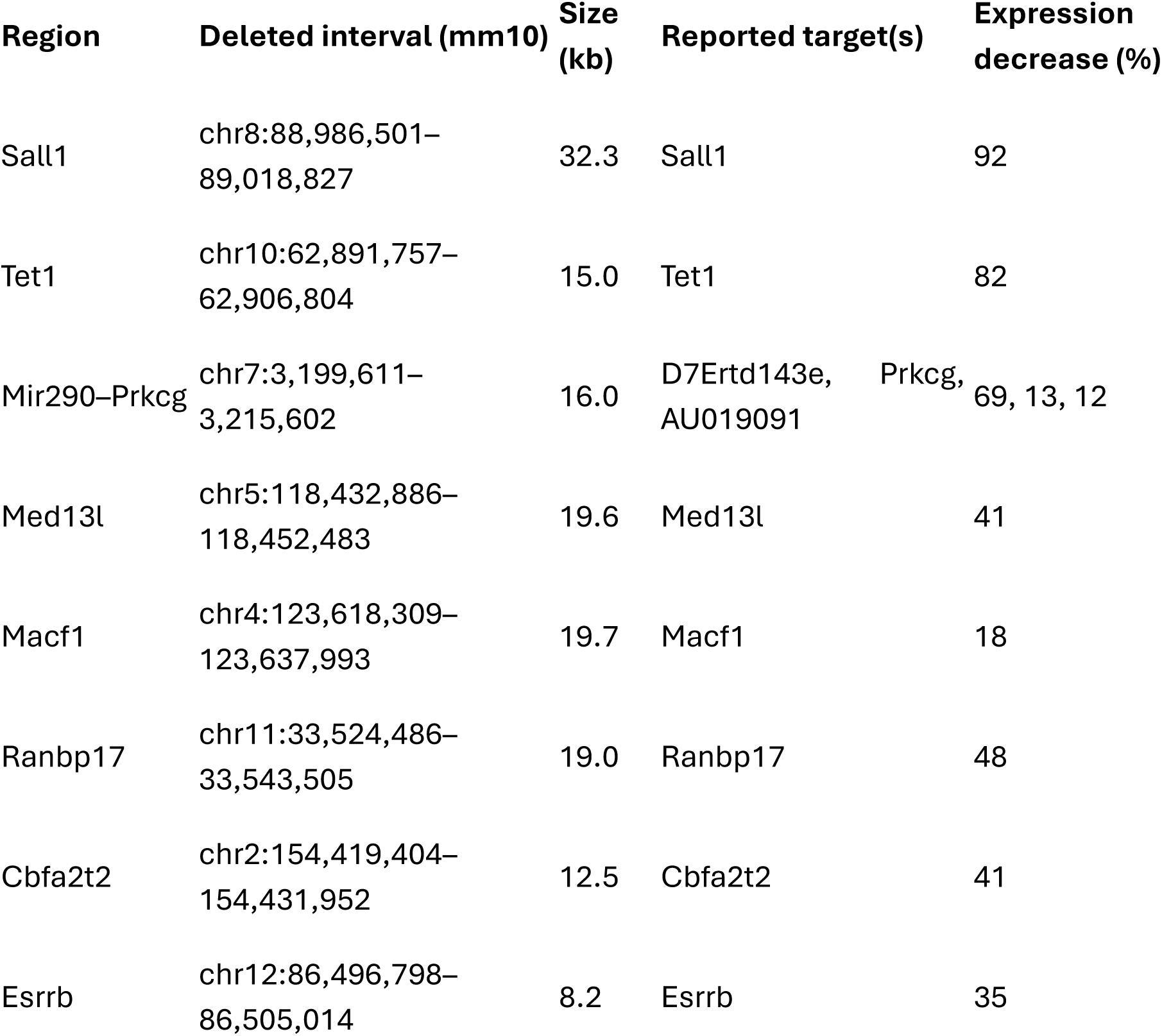
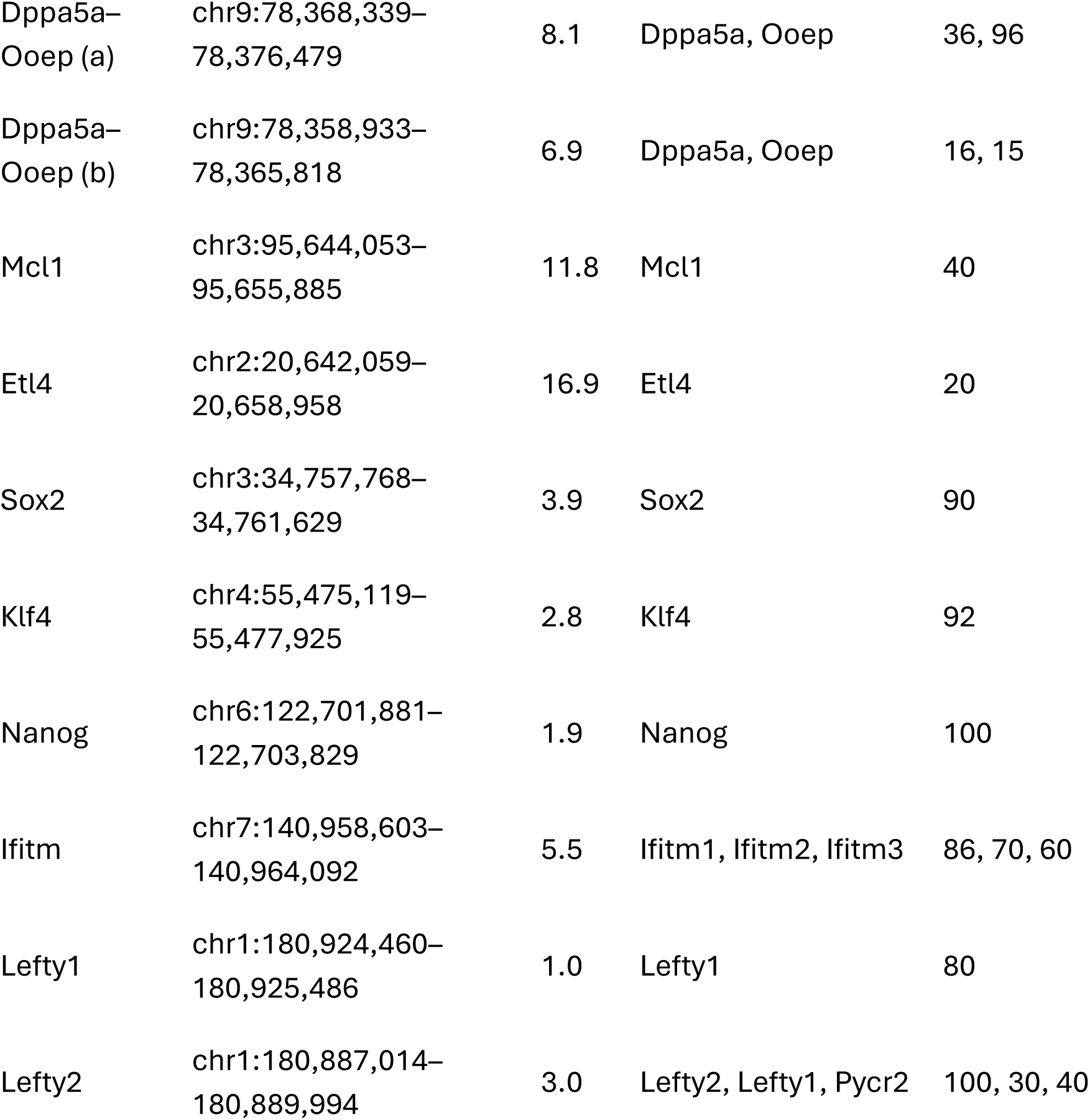
The 18 mESC enhancer deletions. Coordinates are mm10. Expression decrease is the reported percent loss for each target, in the same order as the targets.

**Table 3.** Model hyperparameters.

| Parameter | Value |
| --- | --- |
| tree_method | hist |
| objective | multi:softprob (6 classes) |
| device | cuda:0 |
| eval_metric | aucpr |
| learning_rate | 0.015 |
| n_estimators | 6000 |
| early_stopping_rounds | 150 |
| max_depth | 8 |
| max_leaves | 63 |
| min_child_weight | 8 |
| gamma | 0.05 |
| reg_alpha | 0.05 |
| reg_lambda | 5.0 |
| subsample | 0.65 |
| colsample_bytree | 0.4 |
| colsample_bylevel | 0.6 |

#### Features

The model used 422 features; features are summarized in Table S4.

### Gini validation

#### CAGE expression data

Promoter-level expression was taken from the FANTOM5 phase 1 and 2 mouse CAGE peak atlas (mm10_fair+new_CAGE_peaks_phase1and2_tpm.osc.txt)(ref *55*). It contains 164,672 CAGE peaks across 1,073 samples, with expression given as RLE-normalized tags per million (TPM). Peaks with mm9 coordinates were converted to mm10 using the UCSC mm9ToMm10 liftOver chain. We kept peaks with TPM > 1 in at least 10 samples. This left 133,639 expressed peaks, and all analyses below use this set.

#### Promoter expression statistics

We summarized each peak’s expression across the 1,073 samples with three statistics. The Gini coefficient was computed on log(TPM + 1) values. Dynamic range was the difference between the 95th and 5th percentiles of log(TPM + 1).

#### Peak-to-gene assignment

Peaks were assigned to genes using a TSS annotation derived from the UCSC mm10 NCBI RefSeq gene models, in which TSSs within 300 bp of one another were merged. This gave 69,806 TSSs for 49,997 genes. Each peak was assigned to the nearest TSS within 200 bp of the peak midpoint, without regard to strand. Where several peaks were assigned to one gene, we kept the peak with the highest mean log(TPM + 1). This gave one promoter for each of 22,871 genes, of which 20,314 were in the expressed set; we refer to these as annotated TSSs. The remaining 113,325 expressed peaks (85%) were not assigned to a gene.

#### Housekeeping and developmental gene sets

Housekeeping genes were the 3,277 mouse genes in the Housekeeping and Reference Transcript Atlas v1.0 (HRT Atlas) Hounkpe et al., 2021 (*63*). That atlas defines housekeeping genes by stable expression across 507 RNA-seq samples from 14 C57BL/6 tissues and cell types. Developmental genes were the 512 genes bound by all four Polycomb group proteins (Suz12, Eed, Phc1 and Rnf2) within 1 kb of the TSS in mouse ES cells, as measured by ChIP-chip Boyer et al., 2006 (*64*). This set is defined by Polycomb binding and is enriched for developmental transcription factors. HRT Atlas membership is defined by expression stability, so it is not independent of the breadth and Gini statistics. The Boyer set is defined independently of expression.

Gene names from each list were first matched directly to the gene names of the annotated TSSs. Names that did not match were resolved through NCBI Gene (Mus_musculus.gene_info): by Entrez Gene ID for the Boyer set, and by symbol or synonym for the HRT set. A match was kept only if it resolved to exactly one NCBI gene and exactly one annotated gene. This matched 3,172 of 3,277 HRT genes and 459 of 512 Boyer genes. Unmatched genes either had no CAGE peak within 200 bp of their TSS or had an ambiguous alias. Eight genes appeared on both lists and were removed from both. The final expressed set contained 3,144 housekeeping and 443 developmental promoters, one per gene.

#### UMAP embedding

For the embedding, each peak’s log(TPM + 1) profile was z-scored across samples, so that the embedding reflects expression pattern rather than level. We drew 50,000 expressed peaks at random (seed 42) and embedded them with UMAP using 30 neighbours, a minimum distance of 0.3, cosine distance and random seed 42. The random sample included 1,188 housekeeping and 157 developmental promoters (*67*).

#### Distributions

Distributions of Gini and dynamic range were estimated with Gaussian kernel density estimates (KDEs), with bandwidth set by Scott’s rule. For all expressed peaks, the KDE was fitted to a random sample of 20,000 peaks. In the joint Gini × dynamic range plot, each class’s 50% and 90% contours were drawn at the density levels corresponding to the 50th and 10th percentiles of that class’s KDE evaluated at its own points.

